# High-salinity sporulation in *Bacillus subtilis* results in coat dependant enhanced resistance to both wet heat and hydrogen peroxide

**DOI:** 10.1101/2025.06.24.660303

**Authors:** Víctor Freire, Irene Orera, Santiago Condón

## Abstract

In natural niches such as soil, spore-formers encounter fluctuating conditions, leading to sporulation in non-optimal environmental conditions that can affect spore germination and resistance properties. However, while the influence of sporulation temperature on spore behaviour has been widely studied, the data concerning the influence of other factors such as water activity (*a_w_*) is scarce. We reported that reducing the *a_w_*of the sporulation medium from the optimum (*a_w_* 0.99; S_control_ spores) to 0.98, using sodium chloride (S_salt_ spores) increased their wet heat resistance, yet by unknown means. The present work aimed to examine the mechanism behind the increased wet heat resistance observed in the spores produced in the presence of high salinity. Crust morphogenetic protein CotY was required for the increased wet heat resistance observed in spores of *Bacillus subtilis* formed in the presence of salt. Interestingly, this protein was also related to increased H_2_O_2_ resistance. Proteomic analysis of coat extracts from S_salt_ and S_control_ spores revealed markedly different protein profiles.

Specifically, S_salt_ spores displayed significant increase in abundance of proteins involved in redox homeostasis, structure of the coat and also, others involved in favouring or establish cross-linking. Increased cross-linking, such as disulfide bond formation in the coat and the cysteine-rich crust is known to influence resistance to both agents. Furthermore, deletion of CotY increased DPH lipid probe mobility regardless of sporulation condition, indicating that it plays an active role in stabilizing the IM. Therefore, all the presented data suggested that increased coat cross-link directly or indirectly related to CotY in S_salt_ grants enhanced wet heat resistance by protecting the IM, which was supported by DPA release data under lethal heat stress. This work advances our understanding of how the coat modulates resistance in bacterial spores, helping develop effective control strategies against problematic spore populations due to soil salinization.

**Graphical abstract:** 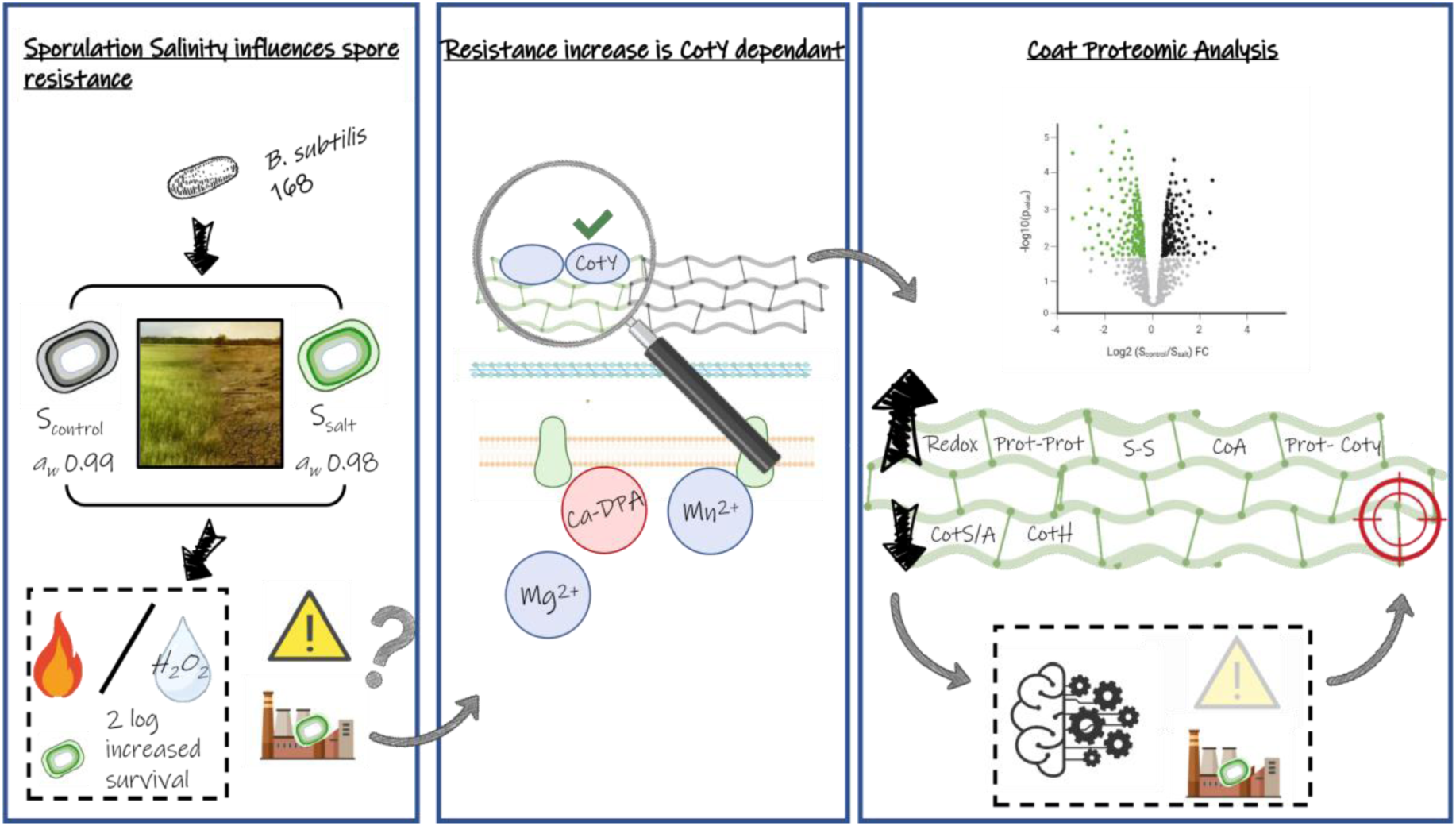

## Introduction

The extraordinary resistance of bacterial spores enables them to persist and disperse in the environment over long times and distances (Chen et al., 2010; Enger et al., 2018; Wijman et al., 2007). This makes them ubiquitous, rendering their entrance into the food chain almost unavoidable (Beskrovnaya et al., 2021). Once there, these spores withstand all but the most intense sterilization treatments applied to foods. Survivors may then germinate in response to the nutrients present (André et al., 2017). Subsequently, germinated spores outgrow into vegetative cells, regaining the ability to multiply and produce toxins, ultimately compromising food stability and safety (André et al., 2017; Setlow & Johnson, 2019). While increasing the intensity of heat treatments may seem like a straightforward solution to mitigate these risks, thermal sterilization negatively impacts the organoleptic and nutritional qualities of foods (Sevenich & Mathys, 2018).

These facts have justified decades of research into the resistance mechanisms of bacterial spores against various agents, especially wet heat - the principal spore inactivation method used in the food industry - with the aim of developing alternative, cost-effective processes that better preserve quality. As a result, these investigations have revealed that this extraordinary resistance is granted by a unique structure and composition, more precisely, by superimposed layers that engulf a dehydrated core delimited by the inner membrane (IM). This dehydration is one of the fundamental factors explaining the enormous difference in wet heat resistance compared to vegetative cells of the same species, several orders of magnitude apart (Setlow, 2014). During sporulation, water content is partially replaced by minerals, with one particular form of calcium and dipicolinic acid chelate (Ca-DPA) comprising around 25 % of the dry weight of the spores and contributing significantly to its resistance against wet heat (Kanaan et al., 2022; Setlow et al., 2006). Furthermore, the properties of the inner membrane (IM) were also found to significantly contribute to resistance against wet heat (Berendsen, Boekhorst, et al., 2016; Berendsen, Koning, et al., 2016; Flores et al., 2023; Kanaan et al., 2022; Korza et al., 2023).

Core dehydration is maintained by the cortex, a modified peptidoglycan layer which compresses the IM. Likewise, the cortex is surrounded by several layers of proteinaceous nature, the coat, which has the function to regulate entry of molecules, such as germinants, repel or neutralize chemicals and resist phagocytosis (Ghosh et al., 2018; Kanaan et al., 2022; Klobutcher et al., 2006; Malyshev et al., 2021; Saggese et al., 2022). These layers vary among spore-formers, in *B. subtilis* the coat is composed of the cysteine-rich outermost layer the “crust”, the outer coat, inner coat, and basement layer, adjacent to the cortex (McKenney et al., 2013). Importantly, this complex coat structure and the degree of cross-linking were confirmed to also significantly influence resistance against wet heat, and were hypothesized to be a consequence of IM stabilization during heat stress (Abhyankar et al., 2015; Kanaan et al., 2022).

It is important to note that, many of these structures and properties related to wet heat resistance can be altered by the environmental conditions in which spores are formed. For instance, modifications have been reported in the water core content, inner membrane, and also coat composition (Beaman & Gerhardt, 1986; Bressuire-Isoard et al., 2016; Cortezzo & Setlow, 2005; Isticato et al., 2020). The influence of these environmental conditions on wet heat resistance has been studied in detail in the case of sporulation temperature (Bressuire-Isoard et al., 2018; Huang et al., 2023; Melly et al., 2002), yet it has been neglected in the case of conditions such as water activity (*a_w_*). With only some studies covering this topic in *B. subtilis* (Nguyen Thi Minh et al., 2008, 2011), showing a decrease in the resistance of spores formed at *a_w_* 0.95. On the other hand, a previous characterization of the influence of different environmental conditions on resistance of spores formed under equally stressful conditions, found an increase in the wet heat resistance of bacterial spores obtained at *a_w_* 0.98 with either glycerol or NaCl (Freire et al., 2023). As to date, design of treatments aiming to inactivate spores to guarantee a degree of stability and innocuity is made according to data obtained from inactivation studies of spores obtained under optimal sporulation conditions (Bressuire-Isoard et al., 2018). However, the findings presented in several studies point towards the need to incorporate environmental influence into this, as conditions in the natural sporulation niches are subject to constant change and could greatly alter resistance of bacterial spores more than 10-fold (Freire et al., 2023). This is especially true for conditions such as increased temperature and higher salinization, which are expected to intensify due to global warming and its impact on soil salinization (EEA. 2019; IPCC. 2022).

Due to this, we decided to further investigate the properties of the spores formed under the conditions previously mentioned of high salinization due to their problematic profile. These spores may compel the application of heat treatments of increased intensity if encountered contaminating foods due to their increased heat resistance. Furthermore, combined treatments leveraging on the existence of a significant subpopulation that sustained sublethal damage may not be as effective as in the case of spores formed under optimal conditions as in S_salt_ population the sublethal damage accumulation would be slower (Freire et al., 2023). *B. subtilis* spores frequently contribute to food spoilage, and certain strains can cause food-poisoning incidents via toxin production (André et al., 2017; Apetroaie-Constantin et al., 2009). In addition, spores of this species also serve as a valuable model for studying mechanisms of bacterial-spore inactivation. Therefore, further insight into the mechanisms of this increased resistance will contribute to the development of new procedures to control problematic spore populations on the rise due to environmental changes. The present work achieves to advance in this matter by offering information about the role of the coat in the increased wet heat and also, H_2_O_2_ resistance. Remarkably, Coty crust morphogenetic protein was needed for the development of this increased resistance, likely by stabilization of the IM. Finally, protein composition changes occurring in the coat leading to this increased resistance, are examined and discussed.

## Material and methods

### Obtaining and purifying of spore suspensions

*B. subtilis* 168 was used throughout this study. The strain was maintained at −80 °C in nutrient broth No. 2 (NB; Oxoid, Basingstoke, UK) supplemented with 25 % glycerol. For revitalization, cells were streaked on nutrient agar (Oxoid) supplemented with 0.6 % yeast extract (NAYE; Oxoid) and incubated at 37 °C for 24 h. For sporulation and spore harvest the same protocol as indicated in Freire et al. (2023) was used. In addition to WT strain, in some experiments strains lacking CotY or CotE coat morphogenetic proteins were used. These deletion strains were constructed by SPP1 phage transduction as previously described (Burton et al., 2019), using as donor strains the deletion mutants (BKE11750 and BKE17030) from the BKE genome-scale deletion library (NBRP - National BIO-Resource Project, Japan) developed by Koo et al. (2017).

### Thermal treatment

To examine the inactivation kinetics, heat treatments at 105 °C were carried out in a specially designed thermoresistometer (Condón et al., 1993). Viability and DPA release kinetics were assessed by treating samples at 95 °C in a PCR thermocycler (Bio-Rad T100, Hercules, CA, USA). Released DPA content was determined by terbium fluorometry, using autoclaved samples as the 100% fluorescence reference, as detailed in (Freire et al., 2024).

### Determination of survival

Survival was routinely determined by pour-plating in NAYE. NAYE plates were incubated at 37 °C for 24 h. Longer incubation times did not affect the survival counts. Plate counts were obtained using an automatic colony counting system by image analysis. The survival fraction was calculated as the difference between the logarithm of *N*_t_ and *N*_0_ (log (*N*_t_*/N*_0_)), which represent the number of survivors in CFU/mL after different treatment times and prior to treatment, respectively.

### Calculation of heat resistance parameters

Since the survival curves had shoulders, experimental data were fitted to the Log-linear + shoulder equation propounded by Geeraerd et al. (2000) (Eq. (1)), using the GInaFiT Excel tool (Geeraerd et al., 2005; KU Leuven, Leuven, Belgium). This equation describes survival curves by means of two parameters: shoulder length (*Sl*, min), defined as the time required to reach the exponential inactivation rate, and the inactivation rate (*k_max_*, min^−1^), defined as the slope of the exponential section of the survival curve. The GInaFiT software also provides the R^2^ and root mean square error (RMSE) to assess goodness of fit.

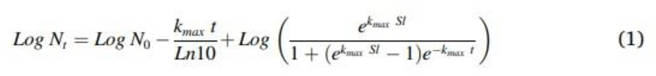

### Resistance to H_2_O_2_

Resistance to H_2_O_2_ was assessed as previously described, with minor modifications (Kanaan et al., 2022; Korza et al., 2023). Briefly, samples were incubated in 10 % H₂O₂ at 25 °C for inactivation, then diluted 1:10 into a catalase solution to neutralize remaining H₂O₂ (Kanaan et al., 2022).

### Coat protein extraction, electrophoresis and Label Free Quantification (LFQ)

This process was realized by boiling the spores in a mixture of reducing agents SDS-DTT with the aim to solubilize the coat proteins as described previously by other authors in more detail (Isticato et al., 2020). The protein content of the different samples was measured by using RC DC assay kit (BIORAD, USA). Equal amounts of protein were loaded into each lane of precast 4-15 % polyacrylamide (BIORAD) gels prior to electrophoresis for initial assessment of differences.

For LFQ, an independent set of identical samples was cleaned in-gel by SDS-PAGE, allowing them to run only until the proteins entered the top of the resolving gel. Subsequently, a single band was excised for each sample with a scalpel in a biosafety cabinet to avoid contamination with keratins. The digestion of excised samples was carried out in an automatic digester (Intavis, Bioanalytical Instruments, Cologne, Germany). Briefly, the bands were washed with water, ammonium bicarbonate (100 mM), and acetonitrile (ACN). The samples were then reduced by incubation with DTT (10 mM) at 60 °C for 45 min and alkylated by incubation with iodoacetamide (50 mM) at room temperature for 30 min in the dark. Finally, proteins were digested with trypsin overnight at 37 °C (5 ng/μl, Trypsin Gold, Promega, WI, USA). The digestion was stopped by adding 0.5 % TFA (trifluoroacetic acid), and the tryptic peptides were extracted sequentially with increasing concentrations of ACN in water.

Proteins were identified and quantified using a hybrid trapped ion-mobility quadrupole timeof-flight mass spectrometer (TIMS TOF Flex, Bruker Daltonics, Bremen, Germany) coupled online to an EvoSep ONE liquid chromatograph (EvoSep, Odense C, Denmark). Peptide quantification was carried out using the Qubit kit according to the manufacturer’s instructions. A total of 200 ng of digested samples were directly loaded onto the EvoSep ONE chromatograph, and profiles were acquired using the 60 SPD (samples per day) protocol.

Peptides were separated on a C18 column (8 cm × 150 μm, 1.5 μm, Evosep) using a linear 21minute gradient and a cycle time of 24 minutes at a constant flow rate of 1 μL/min. Column temperature was controlled at 40 °C. Data were acquired using data-dependent acquisition mode with PASEF (parallel accumulation serial fragmentation). MS data were collected over an m/z range of 100 to 1700 and a mobility range of 0.60–1.60 V·s/cm². During each MS/MS data collection, each TIMS cycle lasted 1.1 seconds and included one MS and ten PASEF MS/MS scans.

Protein identifications and quantifications were performed using Bruker ProteoScape™ (BPS) software and searched against the UniProt *Bacillus subtilis* protein database plus sequences of known contaminants. A precursor tolerance of 20 ppm and a fragment ion tolerance of 30 ppm were used. The search space included all fully- and half-tryptic peptide candidates with up to two missed cleavages. Carbamidomethylation (+57.02146) of cysteine was considered a static modification. TIMScore was enabled to incorporate peptide Collisional Cross Section (CCS) during the scoring process. Identifications were rescored using an updated version of MS2Rescore. These results were validated, assembled, and filtered using a false discovery rate (FDR) of 0.01; under these filtering conditions, the estimated FDR was below ∼1 % at the protein level in all analyses.

Data were filtered to remove proteins with fewer than two peptides, as well as contaminants and reverse hits. The cleaned, quantified protein data file was statistically analyzed and visualized using Perseus (version 2.0.3.0) (Tyanova et al., 2016). Five biological replicates obtained under different working days were included for the LFQ assays.

### Statistical analysis

Statistical analyses (Student’s *t-*test) of experiments other than LFQ were performed using GraphPad PRISM 5.0 (GraphPad Software Inc., San Diego, CA, USA), and differences were regarded as significant when *P* was ≤ 0.05. Data depicted in figures and tables, correspond to averages and standard deviations calculated from at least three biological replicates.

## Results

### S_salt_ increased resistance is not due to increased DPA content per spore, nor higher wet density

The main established factor contributing to spore heat resistance is low core water content, partially replaced by dipicolinic acid (DPA), which stabilizes essential structures such as the core and membrane-associated metabolic machinery (Beaman et al., 1982; Kanaan et al., 2022; Setlow et al., 2006). Given that water and DPA content are known to change with environmental sporulation conditions (Beaman & Gerhardt, 1986; Bressuire-Isoard et al., 2018), suspensions obtained under both optimal (*S*_control_) and high-salinity (*S*_salt_) sporulation conditions were examined for changes in DPA content and wet density. In this regard, no significant differences (*P* > 0.05) were found in the amount of dipicolinic acid content per spore, with around 0.8 ± 0.08 picograms of DPA per spore, regardless of sporulation condition. Similarly, no significant differences arose among sporulation conditions for wet density, with density values of around 1.3 g/cm^3^ (Fig. 1). The content of DPA and the wet heat density observed is in accordance to data reported for *B. subtilis* spores obtained under optimal sporulation conditions in the literature (Jamroskovic et al., 2016; Rose et al., 2007). These results indicated that the increase in wet heat resistance exhibited by S_salt_ spores was unlikely to be due to lower water content or increased Ca-DPA mediated stabilization during heat stress.

**Figure 1:**
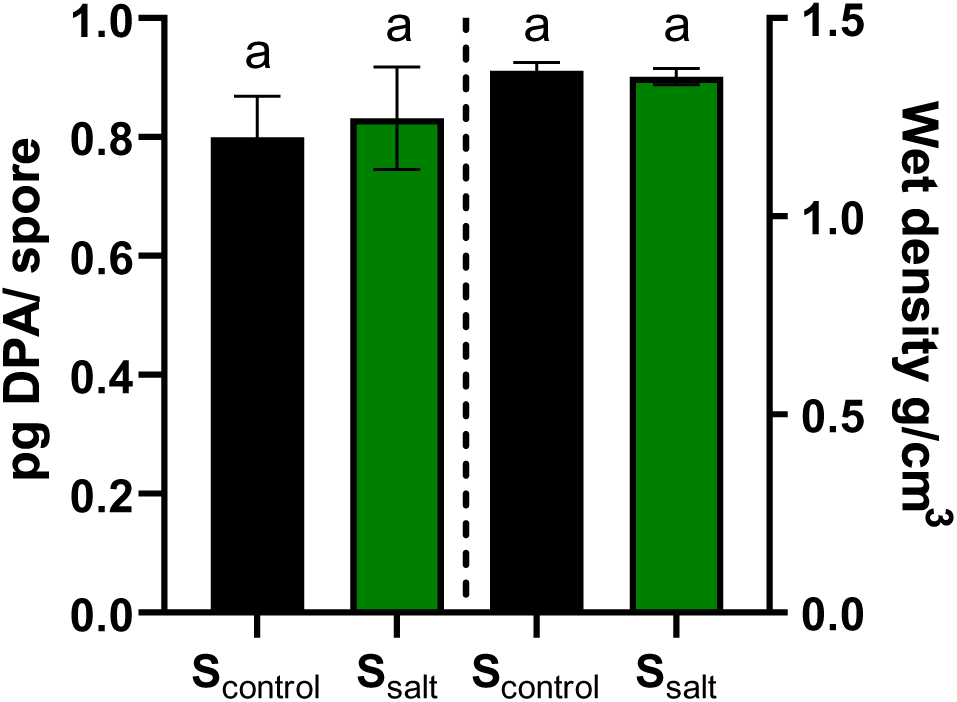
DPA content per spore and wet density of *B. subtilis* spores obtained under optimal (S_control_, black bars) and high salinity conditions (S_salt_, green bars). Different letters indicate statistically significant differences (*P* ≤ 0.05) among sporulation conditions for each property assayed.

### Crust morphogenetic protein CotY contributes to the increased wet heat resistance of spores formed in presence of salt

It has been shown that both the coat structure itself and a decrease in lipid mobility of the IM can substantially increase wet heat resistance in spores of this species (Kanaan et al., 2022; Korza et al., 2023). In addition, both structures have been associated with differences in the inactivation kinetics when exposed to various agents, such as sodium hypochlorite or dodecylamine and even alternative preservation technologies such as UV-C. However, UV-C resistance appears to be more specifically linked to coat modifications (Clair et al., 2020). Notably, S_salt_ spores were more sensitive to sodium hypochlorite and UV-C (Freire et al., 2023), as well as dodecylamine (Freire et al., 2025). In this sense, sporulation of *Bacillus subtilis* 168 at high salinity was found to influence nutrient germination and coat permeability (Freire et al., 2025).

Given the evidence of coat structural modifications under these sporulation conditions, we evaluated whether this structure also contributed to the increase in wet heat resistance. For this purpose, WT and coat-deficient mutant spores were sporulated under both optimal and high-salinity conditions. The Δ*cotY* mutant, lacking the main structural component of the crust, fails to form a complete crust layer (Bartels et al., 2019; Freire et al., 2025). Meanwhile, the Δ*cotE* mutant produces spores in which the outer coat and the crust structures are absent (Chada et al., 2003; McKenney et al., 2013). Subsequently, survival curves at 105 °C were obtained in McIlvaine buffer of pH 7.0 and spores recovered under optimal conditions (Fig. 2).

**Figure 2:**
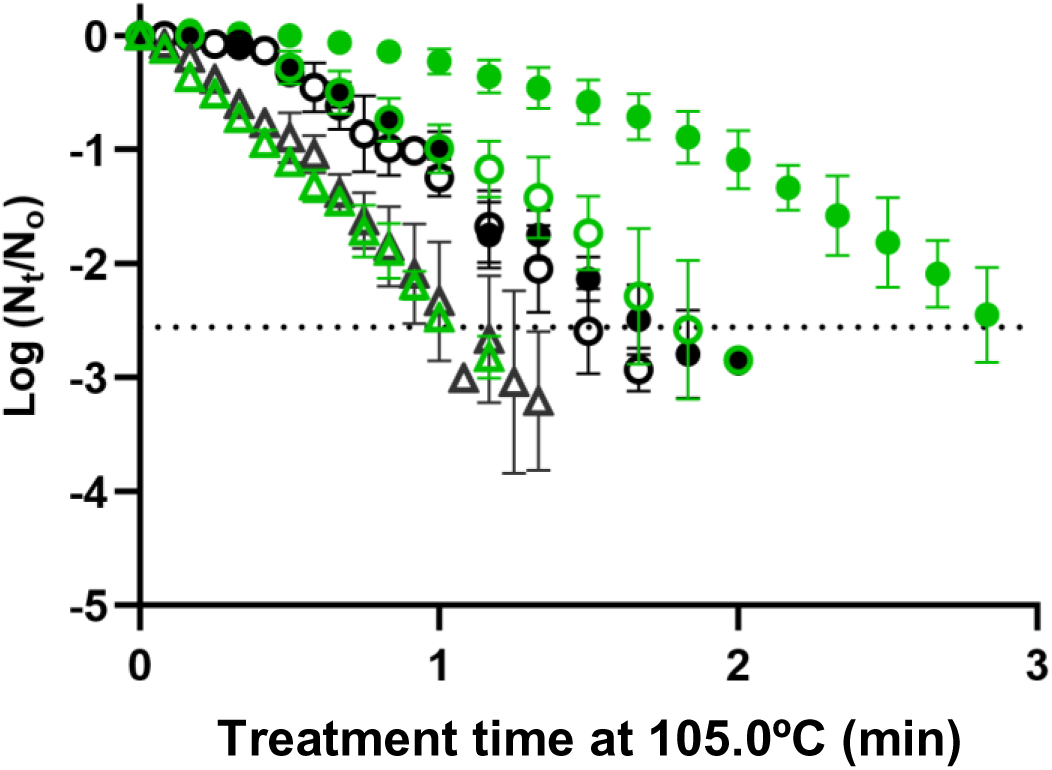
Survival curves at 105.0 ◦C of spore populations of *B. subtilis* under optimal (black) and high salinity conditions (green), WT spores are depicted with full symbols 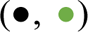, whilst coat defective strains are depicted by empty symbols; Δ*cotY* (○, ○) and Δ*cotE* (Δ, Δ), respectively.

### Treatment time at 105.0°C (min)

Finally, their inactivation kinetics were modelled (Table 1). S_salt_ WT spores presented a higher wet heat resistance, characterized by an increased shoulder length, lower inactivation rate (*k_max_*) and a higher 3D_105 °C_ value in comparison to S_control_ WT spores, as previously described in more detail (Table 1, P ≤ 0.05, Freire et al., 2023).

**Table 1:**
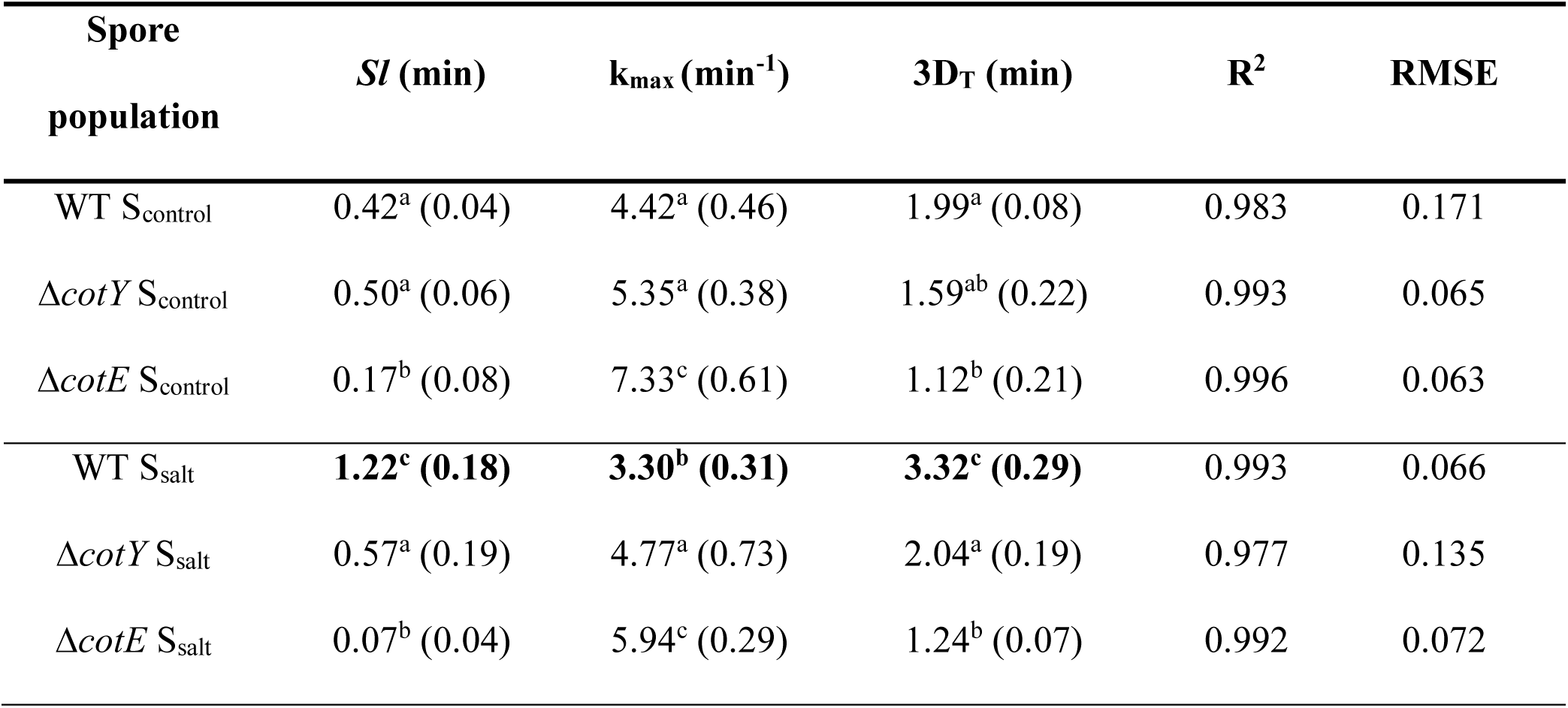
Heat resistance parameters of the different spore populations obtained from the fit of the Log-lineal + shoulder model (Geeraerd et al., 2000) to survival curves at 105 °C (Fig.2). Survivors were recovered in NAYE. Data in brackets represent the standard deviations of the means. Bold letters indicate statistically significant differences (*P* ≤ 0.05) in resistance parameters among sporulation conditions within each strain. Lowercase letters indicate statistically significant differences among suspensions (*P* ≤ 0.05).

| Spore population | *Sl* (min) | $k_{\max}$ (min<sup>-1</sup>) | 3D<sub>T</sub> (min) | R<sup>2</sup> | RMSE |
| --- | --- | --- | --- | --- | --- |
| WT *S*<sub>control</sub> | 0.42<sup>a</sup> (0.04) | 4.42<sup>a</sup> (0.46) | 1.99<sup>a</sup> (0.08) | 0.983 | 0.171 |
| $\Delta cotY$ *S*<sub>control</sub> | 0.50<sup>a</sup> (0.06) | 5.35<sup>a</sup> (0.38) | 1.59<sup>ab</sup> (0.22) | 0.993 | 0.065 |
| $\Delta cotE$ *S*<sub>control</sub> | 0.17<sup>b</sup> (0.08) | 7.33<sup>c</sup> (0.61) | 1.12<sup>b</sup> (0.21) | 0.996 | 0.063 |
| WT *S*<sub>salt</sub> | **1.22<sup>c</sup> (0.18)** | **3.30<sup>b</sup> (0.31)** | **3.32<sup>c</sup> (0.29)** | 0.993 | 0.066 |
| $\Delta cotY$ *S*<sub>salt</sub> | 0.57<sup>a</sup> (0.19) | 4.77<sup>a</sup> (0.73) | 2.04<sup>a</sup> (0.19) | 0.977 | 0.135 |
| $\Delta cotE$ *S*<sub>salt</sub> | 0.07<sup>b</sup> (0.04) | 5.94<sup>c</sup> (0.29) | 1.24<sup>b</sup> (0.07) | 0.992 | 0.072 |

Conversely, inactivation parameters of S_control_ Δ*cotY* and S_salt_ Δ*cotY* spores showed no statistically significant differences among them nor when compared to S_control_ WT spores (*P* ≤ 0.05; Table 1; Fig. 2). Similarly, other authors found that this protein may not influence heat resistance of spores produced under optimal conditions (Zhang et al., 1993). On the other hand, 3D_105 °C_ values of Δc*otE* strain were significantly inferior to those values observed for WT or Δ*cotY* spores, while inactivation parameters were identical among sporulation conditions (*P* > 0.05, Table 1).

Interestingly, even though a higher inactivation rate was appreciated, these differences were mainly caused by the reduction in the shoulder length which was around 4-fold lower in Δc*otE* spores than that of S_control_ WT spores (*P* ≤ 0.05, Table 1). These results demonstrated the relevance of the coat in the wet heat resistance of these spores. This is in agreement with results obtained using other strains and experimental setups, as removal of a substantial part of the coat structure either via mutants lacking morphogenetic proteins or by chemical treatment has also been associated with significant differences in heat inactivation kinetics (Ghosh et al., 2008; Kanaan et al., 2022). Remarkably, the fact that enhanced resistance in S_salt_ spores required CotY, pointed to a modification within the coat in which CotY was involved in as the ultimate responsible of this increase in resistance experienced by S_salt_ WT spores.

### Basal Membrane rigidity would not be the cause for the increased wet heat resistance of spores cultured at high salinity

Since the coat structure and the inner membrane mobility may alter simultaneously the resistance to wet heat (Kanaan et al., 2022; Korza et al., 2023), DPH anisotropy measurements were conducted in order to discard synergistic contribution to the increased wet heat resistance of S_salt_ spores. These measurements offer information about the mechanical behaviour of the spore membranes (Voss & Montville, 2014), and were conducted on spores obtained under both sporulation conditions on both WT suspensions and on Δ*cotY* spores given the important role observed for this protein in the inactivation kinetics.

The values for rigidity were about 0.130 for the case of S_control_ WT spores and S_salt_ WT spores while in the case of S_control_ Δ*cotY* and S_salt_ Δ*cotY* values were around 0.090, thus, no statistically significant differences were observed among sporulation conditions (Fig. 3). These values are on the range observed by other authors; however, it is important to remark that, differences in these values can arise from the species, strain, experimental design and fundamentally from the sporulation medium composition, in addition to the measurement temperature used. For example, anisotropy of spores of *B. subtilis*, *B. anthracis* and *B. mycoides* measured at 30 °C resulted in values around 0.3 to 0.15 anisotropy (Liu, 2014). More importantly, differences were statistically different for WT and Δ*cotY* spores (*P* ≤ 0.05, Fig. 3). This indicated a higher lipid probe mobility in the mutant spores lacking the crust morphogenetic protein CotY.

**Figure 3:**
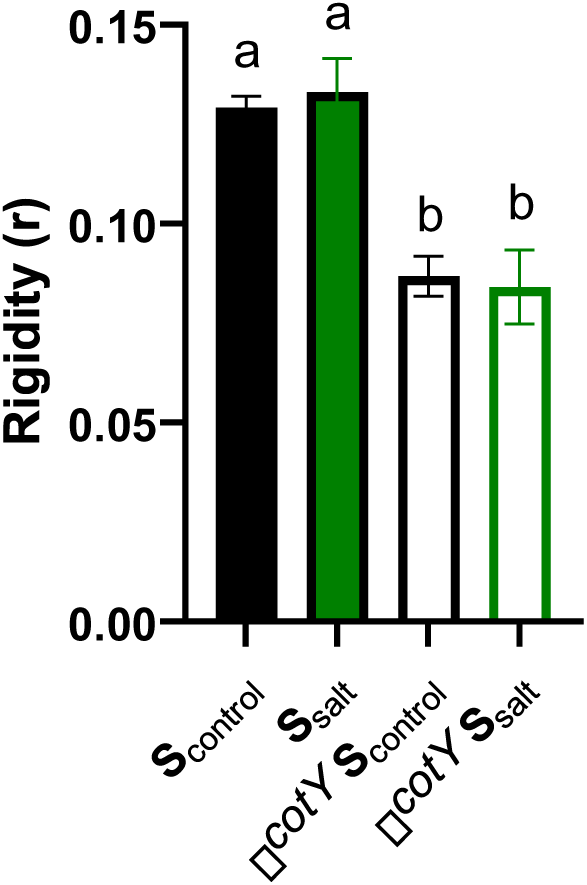
Membrane rigidity of *B. subtilis* spores obtained under optimal (S_control_, black bars) and high salinity conditions (S_salt_, green bars). WT spores are depicted with full bars 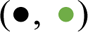, whilst Δ*cotY* spores are depicted by empty bars; (○, ○). Anisotropy measurements were conducted at 37°C.

These results would imply that sporulation at low a_w_ using NaCl as depressor would not result in an apparent change in membrane fluidity, however, CotY would actually play a role in maintaining the low mobility of lipids in the membranes of bacterial spores.

### DPA release in response to heat stress is delayed in spores cultured at increased salinity

In order to study whether or not this coat modification stabilized in some fashion the IM during heat treatments at lethal temperatures, we conducted viability assays at 95 °C and calculated the amount of DPA released by each population of WT and Δ*cotY* spores formed under optimal and high salinity conditions (Fig. 4). Once more, increased wet heat sensitivity was observed in S_control_ spores when compared to S_salt_ WT spores while deletion of CotY eliminated these differences among sporulation conditions (data not shown). Remarkably, a slower DPA release was observed in WT S_salt_ spores, at 45 min this population had released slightly less than 40 % of its total DPA content, which is significantly less (*P* < 0.05; Fig. 4) than that released by WT S_control_ spores with around 59 % DPA loss during this treatment time. In addition, Δ*cotY* spores regardless of sporulation condition liberated around 53 % during the same treatment time, showing no statistical differences with the content released by S_control_ WT spores (*P* > 0.05; Fig. 4). This level of DPA released are similar to those observed by Tu et al. (2021) after heating PY59 spores at 98 °C for 1 hour. DPA release under these conditions is attributed to IM loss of integrity after death of bacterial spores by wet heat (Coleman et al., 2007). Therefore, these results indicate that under heat stress, the integrity of the IM in S_salt_ spores is better preserved by coat modifications which require CotY to take place.

**Figure 4:**
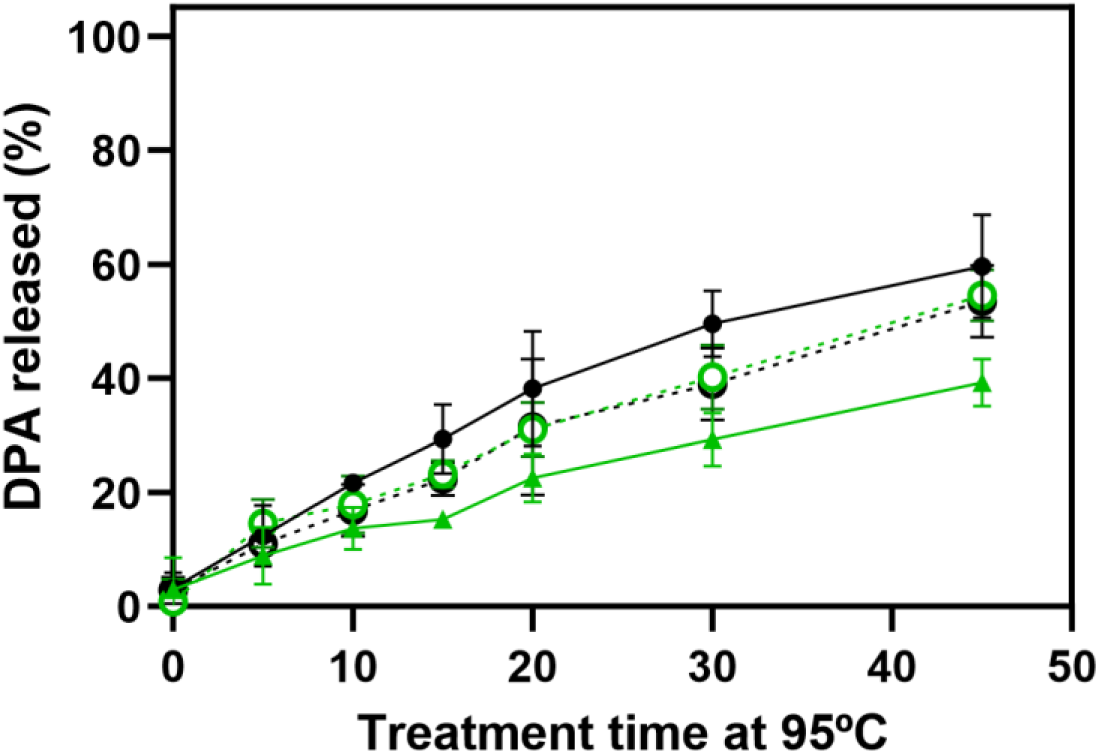
DPA release kinetics of WT and Δ*cotY* spores obtained under both optimal (black symbols) and high salinity conditions (green symbols). WT spores are depicted with full symbols and continuous lines 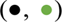, whilst Δ*cotY* spores release are depicted with empty symbols (○, ○) and discontinued lines, respectively.

### H_2_O_2_ resistance is greatly enhanced in S_salt_ spores in a CotY dependant manner

It has been suggested that both wet heat and H_2_O_2_ could kill spores of *B. subtilis* by similar mechanisms, inactivating a key target in the IM (Korza et al., 2023).

In addition, hydrogen peroxide is a disinfection agent widely used in the food industry due to its broad-spectrum antimicrobial activity against bacteria, fungi, viruses, and spores, and finally, ability to breaking down into water and oxygen, leaving no toxic residue (Abdelshafy et al., 2024). Therefore, we decided to characterize the resistance of our suspensions to this agent with the aim to gather further information about the plausible coat modification occurring in S_salt_ spores.

H_2_O_2_ treatment for 20 minutes led to inactivation of ca. 2.3 log cycles of S_control_ WT population, conversely, mean inactivation induced for S_salt_ WT population was significantly less, of ca. 0.3 log cycles (*P* ≤ 0.05, Fig. 5). This indicated that use of this agent against spores formed under high salinity may result in increased risk of survival. On the other hand, S_control_ Δ*cotY* spores were inactivated around 1.8 cycles while S_salt_ Δ*cotY* spores suffered a decrease in the viability of the population of around 1.4 cycles, hence showing no significant differences among them (*P* > 0.05, Fig. 5).

**Figure 5:**
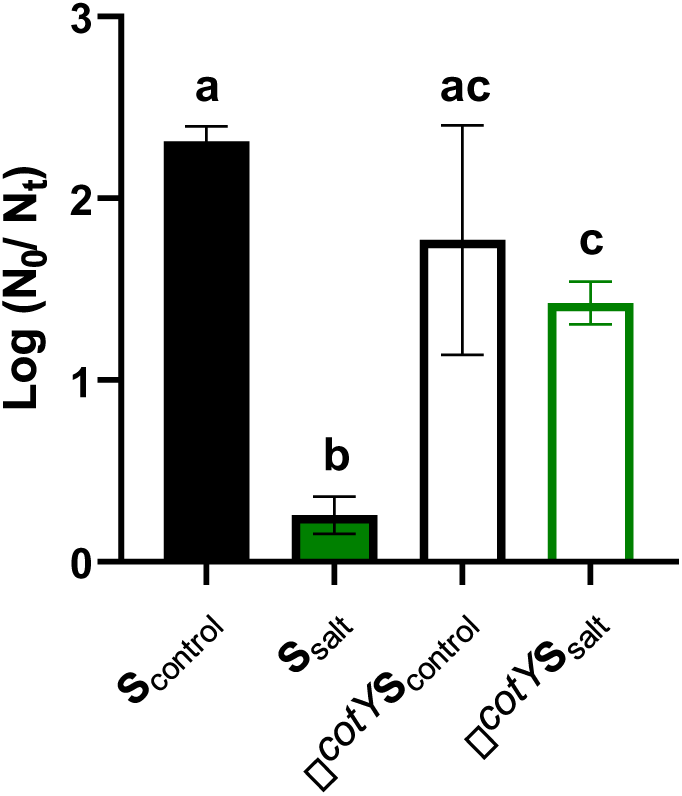
Resistance of S_control_ (Black) and S_salt_ (green) spores (Full bars, WT and empty bars Δ*cotY)* against 20 % H_2_O_2_.

Precisely, the fact that inactivation of S_control_ Δ*cotY* suspensions was not statistically different from that of S_control_ WT spores, while, conversely, spores obtained at high salinity were statistically more sensitive to H_2_O_2_ in spores lacking a complete crust (S_salt_ Δ*cotY*) suggests that a significant modification of the coat, dependent of CotY in S_salt_ spores granted these spores a considerable resistance increase to H_2_O_2_ as well. However, as S_salt_ Δ*cotY* still maintained increased resistance over WT S_control_ spores, these results suggest that additional modifications, such as increased presence of redox-detoxifying coat proteins may also participate in the increased resistance to H_2_O_2_ (see below and Discussion).

### Spores obtained under high salinity conditions suffer significant changes in coat protein composition

Consequently, we hypothesized that changes in the coat that required CotY were behind the increased resistance of S_salt_ spores. In order to gather further information, we decided to assay protein extraction of the coat by SDS-DTT procedure as previously described (Isticato et al., 2020). The extracts were then visualized in a 2D SDS-PAGE gel to check for differences in the solubilized proteins for each sporulation condition (Fig. S2). Coat protein extraction profiles of S_control_ and S_salt_ WT were markedly different, with the intensity of three bands of sizes around 65 kDa, 50 kDa and around 30 kDa visibly diminished in S_salt_ spores. However, the complex nature of the protein-protein interactions within the coat can cause some proteins to be co-extracted with their interacting partners, making identification by this method unreliable (Isticato et al., 2008). Therefore, we opted to subject the coat extracts to label free identification and quantification proteomic analysis to better determine the changes suffered by the coat during sporulation under high salinity conditions.

The proteomic analysis of the coat, depicted in Fig. 6 showed a major alteration of the coat composition between S_salt_ and S_control_ spores. Interestingly, one of the most prominent changes provoked by sporulation at high salinity was significant increase in the abundance of proteins involved in redox homeostasis (Fig. 6, blue dots).

**Figure 6:**
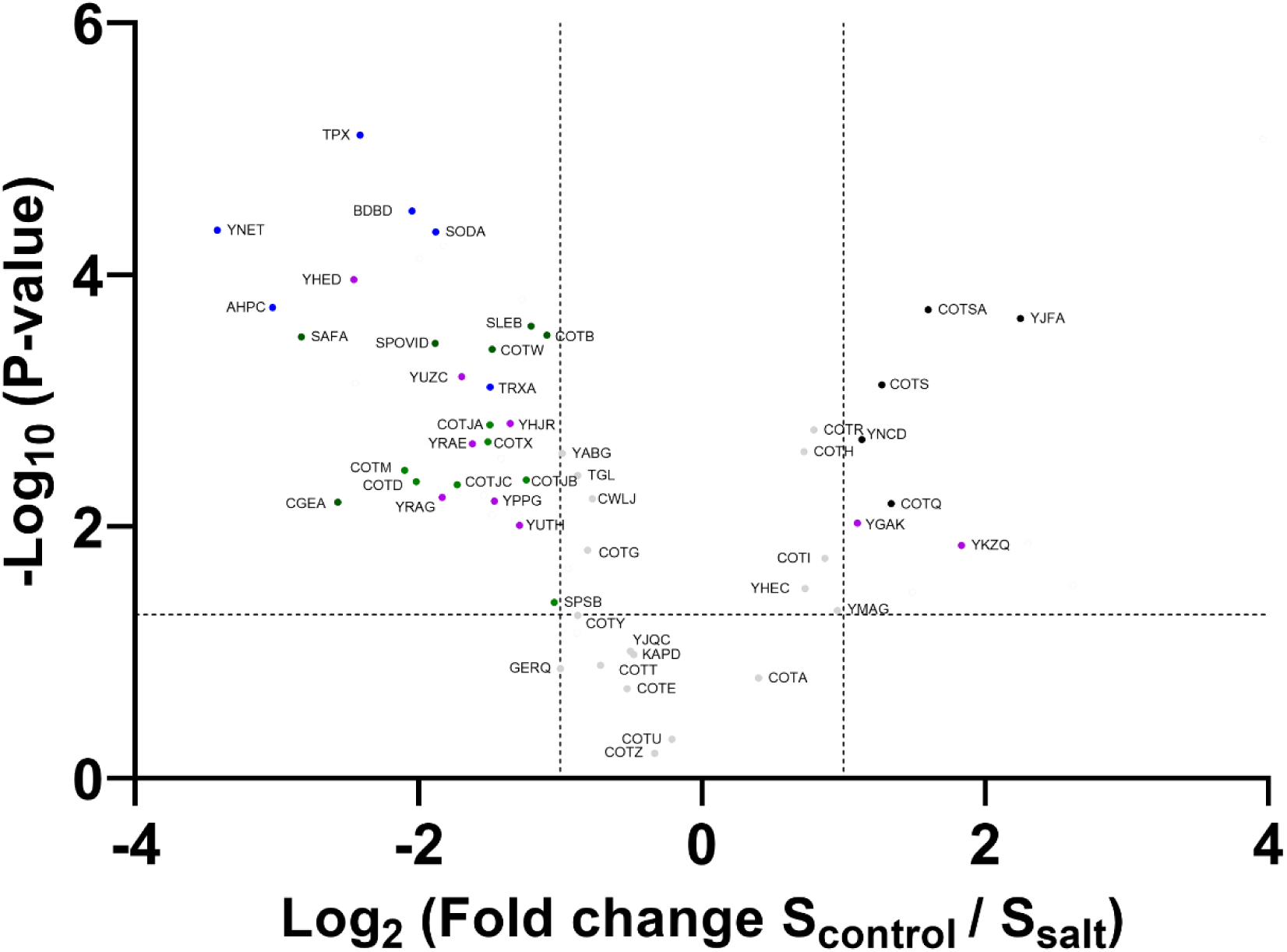
Volcano plot of differentially abundant coat structural or resistance related proteins between S_salt_ and S_control_. Horizontal line delimits the *P* = 0.05 statistically significant level. Vertical lines indicate a Log_2_ Fold Change (FC) in abundance of proteins between both populations. Grey proteins outside these regions are below the FC and *P*-value cut-off to be considered significant. Blue proteins are proteins involved in redox homeostasis and disulfide bond formation. Dark green dots are known structural coat proteins of statistical higher abundance in S_salt_. Black dots are known structural coat proteins of statistical higher abundance in S_control_. Purple proteins are coat proteins of unknown function of statistical higher abundance in either population. Depicted data from five biological replicates

The higher abundance of the thiol-peroxidase Tpx, the disulfide bond formation protein DbdB, the superoxide dismutase SodA, and the thioredoxin TrxA suggested that these cells experienced elevated oxidative stress due to the salinity of the sporulation medium. This observation concurs with results obtained by other studies conducted examining *Bacillus* species, observing induction of peroxidases and superoxide dismutases under salinity stress (Goosens et al., 2013; Hassan et al., 2020). In addition, YneT was also found to be significantly increased in S_salt_. Remarkably, this protein was present in *Bacillus subtilis* spores and known to be CoAlated in a *cotE gerE* dependant manner, although its function still remains obscure (Zhyvoloup et al., 2020).

In addition, structural proteins showed increased abundance in the coat extracts of S_salt_ spores (Fig. 6, green dots). Namely, proteins of every coat layer: basement (SpoVID, YheD), inner coat (SafA), outer coat (CotB, CotM, CotD), and crust (CotW, CotX and CgeA) presented higher abundance in spores formed under high salinity conditions. Conversely, coat kinase proteins such as CotSA and CotS would be upregulated in the spores obtained under optimal conditions (Fig. 6, black dots). Therefore, a major change in structural coat components occurs due to sporulation at high salinity. However, no significant increase was observed in the abundance of the crust morphogenetic proteins CotY. Since this protein was necessary for the development of the increased resistance phenotypes caused by sporulation at high salinity, this could point to protein-protein interaction as the cause for this behaviour.

As a consequence, we examined which of these proteins found in the samples were known to interact with themselves or other coat constituents. 16 proteins with such characteristics were significantly more abundant in spores formed under high salinity conditions, while only 1, CotQ was found more abundantly in spores produced at optimal conditions (Table 2).

**Table 2:**
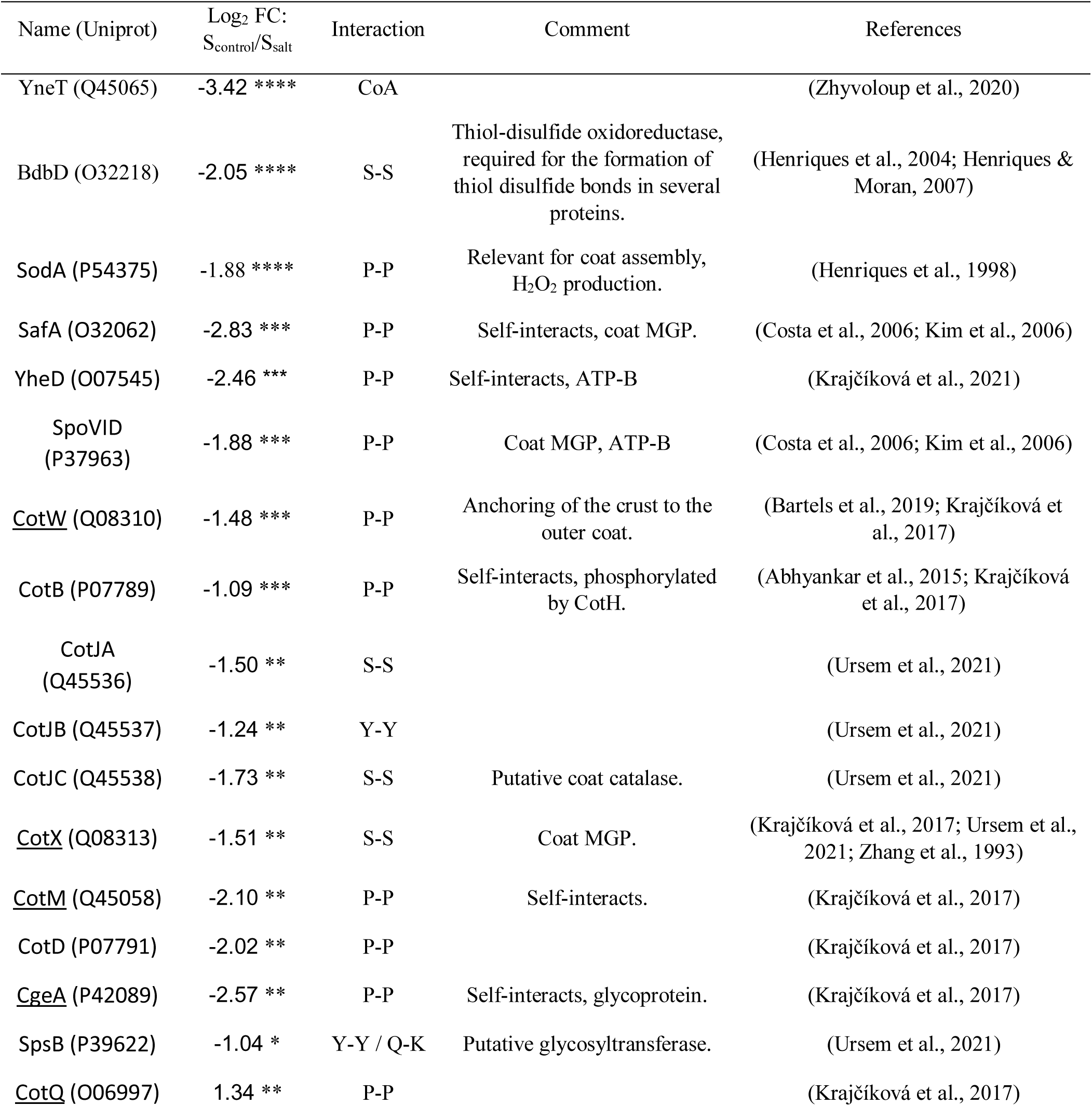
List of coat and coat-related proteins involved in protein-protein interactions. Interaction types: S-S (disulfide), CoA (CoAlation), Y-Y (dityrosine), Q-K (ε-(γ)-glutamyl-lysine), and P-P (undetermined). Statistical significance: *P* < 0.05 (\**), 0.01 (**), 0.001 (\*\*\**) or 0.0001 (****). Data depicted from five biological replicates. MGP: Morphogenetic protein. ATP-B: ATP-binding. Underlined proteins have been experimentally shown to interact directly with CotY.

| Name (Uniprot) | Log<sub>2</sub> FC:<br>S<sub>control</sub>/S<sub>salt</sub> | Interaction | Comment | References |
| --- | --- | --- | --- | --- |
| YneT (Q45065) | -3.42 **** | CoA |  | (Zhyvoloup et al., 2020) |
| BdbD (O32218) | -2.05 **** | S-S | Thiol-disulfide oxidoreductase, required for the formation of thiol disulfide bonds in several proteins. | (Henriques et al., 2004; Henriques & Moran, 2007) |
| SodA (P54375) | -1.88 **** | P-P | Relevant for coat assembly, H<sub>2</sub>O<sub>2</sub> production. | (Henriques et al., 1998) |
| SafA (O32062) | -2.83 *** | P-P | Self-interacts, coat MGP. | (Costa et al., 2006; Kim et al., 2006) |
| YheD (O07545) | -2.46 *** | P-P | Self-interacts, ATP-B | (Krajčiková et al., 2021) |
| SpoVID<br>(P37963) | -1.88 *** | P-P | Coat MGP, ATP-B | (Costa et al., 2006; Kim et al., 2006) |
| <u>CotW</u> (Q08310) | -1.48 *** | P-P | Anchoring of the crust to the outer coat. | (Bartels et al., 2019; Krajčiková et al., 2017) |
| CotB (P07789) | -1.09 *** | P-P | Self-interacts, phosphorylated by CotH. | (Abhyankar et al., 2015; Krajčiková et al., 2017) |
| CotJA<br>(Q45536) | -1.50 ** | S-S |  | (Ursem et al., 2021) |
| CotJB (Q45537) | -1.24 ** | Y-Y |  | (Ursem et al., 2021) |
| CotJC (Q45538) | -1.73 ** | S-S | Putative coat catalase. | (Ursem et al., 2021) |
| <u>CotX</u> (Q08313) | -1.51 ** | S-S | Coat MGP. | (Krajčiková et al., 2017; Ursem et al., 2021; Zhang et al., 1993) |
| <u>CotM</u> (Q45058) | -2.10 ** | P-P | Self-interacts. | (Krajčiková et al., 2017) |
| CotD (P07791) | -2.02 ** | P-P |  | (Krajčiková et al., 2017) |
| <u>CgeA</u> (P42089) | -2.57 ** | P-P | Self-interacts, glycoprotein. | (Krajčiková et al., 2017) |
| SpsB (P39622) | -1.04 * | Y-Y / Q-K | Putative glycosyltransferase. | (Ursem et al., 2021) |
| <u>CotQ</u> (O06997) | 1.34 ** | P-P |  | (Krajčiková et al., 2017) |

Remarkably, CotY-related proteins with a fold change exceeding 1.5 were detected only in S_salt_ spores. For instance, the upregulated proteins observed in S_salt_ such as BdbD (*Bacillus* disulfide bond D) or SodA are known to favour oxidation of thiol groups, such as those present in cysteine contributing to the formation of disulfide bonds of secreted and, perhaps, in the case of the former enzyme, cross-link of coat proteins, aiding to their correct folding and stability (Crow et al., 2009; Henriques et al., 1998). In this sense, two proteins experimentally demonstrated to stablish disulfide bonds in the coat, CotJA and CotJC (Ursem et al., 2021) presented higher abundance in S_salt_ spores. Furthermore, sporulation in S_salt_ also led to a higher abundance of proteins observed to form dityrosine cross-links, such as CotJB and SpsB (Ursem et al., 2021). Finally, some enzymes involved in maturation of the coat structure and acquisition of definitive germination spore properties due to ε-(γ-glutamyl) lysine bonds, such as TgL and YabG (Fernandes et al., 2018, 2019; Monroe & Setlow, 2006; Ragkousi & Setlow, 2004) were also detected to possess higher abundance in S_salt_ populations although their fold change was slightly inferior to 2-fold.

In conclusion, these results show that redox and structural elements, as well as proteins involved in protein-protein interactions such as the various cross-link types described within the spore coat, were significantly more abundant in S_salt_ spores. Importantly, these also included proteins known to interact directly with CotY.

## Discussion

The effect of sporulation conditions, and specially sporulation temperature is widely recognized as an important factor contributing to variability in spore resistance (BressuireIsoard et al., 2018; Freire et al., 2023). However, the impact of *a_w_* has been overlooked so far, despite the fact that environments with lower humidity than ideal culture media, such as soil, are common sporulation niches (Carlin et al., 2010). Especially relevant is the increase in soil salinization as consequence of the tendency to accumulate this solute in soil due to global warming (EEA, 2019; Khamidov et al., 2022). Available reports indicate that in the Mediterranean region, approximately 25 % of cropland is affected by salinization (Geeson,et al., 2003; Mateo-Sagasta & Burke, 2011). For example, in Spain 3 % of the 3.5 million hectares of irrigated agricultural land, is severely affected, and an additional 15 % is at serious risk (Stolte et al., 2016). Previous research demonstrated that *B. subtilis* populations sporulated at depressed *a_w_* (0.98) by the addition of NaCl (S_salt_) exhibited increased wet heat resistance compared to spores incubated under optimal conditions (*a_w_* ∼ 0.99, S_control_). This increase in the thermal resistance was characterized by an increased shoulder length and lower log-linear inactivation rate, most likely due to a slower damage accumulation (Freire et al., 2023).

Increased resistance against the most predominant food preservation technology, along with the foreseeable increase in prevalence of spores formed under these conditions, prompted us to investigate the mechanisms underlying this enhanced resistance. Understanding these mechanisms is essential for designing more effective control strategies. Among the properties granting spores their enormous wet heat resistance, the degree of dehydration, in combination with the DPA and mineral content of the core, plays a major role (Beaman et al., 1982; Beaman & Gerhardt, 1986; Kanaan et al., 2022; Marquis & Bender, 1985; Palop et al., 1999). However, we found no differences in either DPA content nor wet density in S_salt_ spores when compared to those obtained under optimal conditions. Thus, it was unlikely that sporulation at high salinity increased wet heat resistance through either of these mechanisms.

Alternatively, increased wet heat resistance could still be explained by modifications in other structural components, such as the coat and the IM, which are also known to play a significant role, probably in conjunction (Kanaan et al., 2022; Korza et al., 2023). As a consequence, the heat inactivation kinetics and membrane fluidity of WT and mutant spores lacking morphogenetic crust and coat proteins obtained at both optimal and high salinity conditions were examined. WT spores obtained under high salinity retained their characteristic increase in wet heat resistance compared to S_control_ WT spores. In contrast, the inactivation kinetics of Δ*cotY* spores produced under both sporulation conditions were equal and indistinguishable from those observed for S_control_ WT. In addition, loss of additional coat material occurring in Δ*cotE* spores further decreased spore heat resistance, yet again without showing any difference between sporulation conditions. It has been reported that, even after decoating, differences in wet heat resistance were preserved between spores with substantially decreased IM lipid mobility due to 2Duf and/or spoVA^2mob^ alterations, and WT spores with basal IM lipid mobility (Kanaan et al., 2022; Korza et al., 2023). Then, the absence of differences in inactivation kinetics among spores of coat deficient mutants formed under the different sporulation conditions suggested that substantial decrease in the IM mobility by itself was not responsible for the increased resistance observed in S_salt_ spores. Indeed, the lack of differences in DPH anisotropy measurements between sporulation conditions in both WT and ΔcotY spores ruled out salinity-induced changes in the mechanical behavior of the IM under physiological conditions. In conjunction, these results supported the hypothesis that changes in the coat alone, rather than simultaneous significant alterations in both the IM and the coat structures, were the cause of the increased wet heat resistance exhibited by S_salt_ spores.

In addition, the observation that increased resistance occurred only in S_salt_ spores containing CotY supported the involvement of a modification in which this crust morphogenetic protein was involved. In this context, although no significant differences arose in anisotropy values among sporulation conditions, this technique did reveal significant differences between WT and Δ*cotY* spores, with the lack of this protein resulting in significantly lower rigidity values. This is to the best of our knowledge the first experimental measurement of membrane fluidity in spores presenting disrupted coat layers. These findings indicate a higher degree of lipid mobility in the latter spores at physiological temperatures, consistent with the hypothesis that the coat structure plays a role in stabilizing the inner membrane (Kanaan et al., 2022; Knudsen et al., 2015). However, membrane fluidity is known to vary with the temperature to which spores are exposed, as is also observed in vegetative cells (Voss & Montville, 2014), therefore it is unknown if this scenario would also take place under treatment at lethal temperatures.

Nevertheless, during lethal heat stress at 95 °C, final DPA release, as a consequence of loss of IM integrity (Coleman et al., 2007) was greater in S_control_ WT spores than in S_salt_ WT spores, while in Δ*cotY* spores release kinetics were identical among sporulation conditions, and at the same time undistinguishable to S_control_ WT release (*P* > 0.05). Finally, DPA content of both Δ*cotY* and Δ*cotE* spores, showed no substantial differences when compared to WT spores, regardless of sporulation condition (Fig. S1). As a result, these findings suggested that CotYdependent stabilization of the IM occurs during sporulation under high-salinity conditions.

Similarly, an increase in resistance leading to a difference in 2 log cycles of survival to H_2_O_2_ was also observed for S_salt_ WT spores when compared to S_control_ WT spores, while differences in the inactivation of Δ*cotY* spores among sporulation conditions diminished to the point of no statistical significance. Albeit no effect of the coat on the resistance of spores to H_2_O_2_ was found in some studies (Shin et al., 1994), the most recent ones in *Bacillus subtilis* point towards a relevant role of this structure in the protection against this agent (Korza et al., 2023), in agreement with our results. In this regard, spores of *Bacillus subtilis* PS3394 lacking both the crust and the outer coat due to Δ*cotE* deletion, showed decreased resistance against 1 % hydrogen peroxide (Leggett et al., 2016). Furthermore, chemical decoating also increased the sensitivity of spores to this agent, for example, 40 minutes of exposure to 11 % H_2_O_2_ led to a viability difference of over 3 log units between intact and chemically decoated spores of *Bacillus subtilis* PS533 (Kanaan et al., 2022). Therefore, it seems that regardless of using a high or a low concentration of this agent, the coat offers some protection against its action to preserve spore viability. Most likely, H_2_O_2_ must traverse the coat, cross the cortex and the IM to exert damage, or likely act upon it and its components (Korza et al., 2023).

Remarkably, S_salt_ WT spores showed minimal inactivation by this agent, with only ca. 0.3 log reduction after 20 minutes of incubation. As longer incubation times eventually led to inactivation of S_salt_ WT spores (data not shown), this suggests that the coat protective alteration resulting from sporulation under high salinity was depleted with sufficient exposure time. This could be due to two main reasons, (i) the presumably enhanced detoxifying capability of some of the coat components was, finally exhausted. For example, there are redox enzymes present in the coat, such as laccases, which have been shown to effectively protect spores of *Bacillus subtilis* against this agent (Hullo et al., 2001). However, in presence of sufficient concentrations of oxidative agents, protective enzymes may become overwhelmed, leading to irreversible oxidation and loss of function (Arnao et al., 1990). (ii) A presumably decreased permeability could increase over time as H_2_O_2_ strips enough coat proteins for the rest of the molecules of the chemical to reach and react against the IM and the core leading to inactivation of vital enzymes (Gould & Hitchins, 1963; King & Gould, 1969; Korza et al., 2023). Thus, in this way the coat would act like a “reactive armor”, reacting and neutralizing the biocide before it can penetrate further into the spore at the cost of losing integrity (Driks, 1999).

The question of how actually CotY or CotY related coat components in *B. subtilis* could increase resistance against both wet heat and H_2_O_2_ led us to study the composition of this structure in both spore populations. Indeed, in-depth proteomic analysis revealed that shifting sporulation conditions from optimal to high salinity provoked significant changes in the abundance of proteins involved in redox homeostasis, as well as in coat cross-linking. Furthermore, some other enzymatic components of the spore coat such as kinases, which may influence structure and properties such as germination (Saggese et al., 2022) also showed altered abundancy due to sporulation conditions.

Remarkably, CotY, which was needed for the increased resistance of S_salt_ WT spores, showed no significative differences in its abundance. This pointed to interactions among spore coat components, rather than just a higher amount of CotY in the coat of spores produced under high salinity as the responsible for the increased resistance observed. Among these interactions, a higher degree of cross-linking in the coat due to sporulation conditions, may explain the differences in resistance observed in this work. It is known that cross-linking of coat proteins influences wet heat resistance, for example as it is increased by maturation time (Abhyankar et al., 2015; Sanchez-Salas et al., 2011). In some instances, spores with significative differences in heat resistance due to sporulation conditions have also been reported to show altered coat protein expression and cross-linking degree (Abhyankar et al., 2016). Furthermore, it was observed long time ago that pre-treatment with agents rupturing disulfide bonds made *B. cereus* spores more susceptible to H_2_O_2_ action (Gould & Hitchins, 1963), perhaps due to the fact that the cross-linking “hides” reactive sites from H_2_O_2_. Therefore, this mechanism could affect the resistance to both agents. Remarkably, in other species, cysteine-rich proteins of the outermost coat layers, such as, CdeC was required for heat resistance of *Clostridium difficile* spores and also influenced resistance against phagocytic cells, known to exert its action against pathogens by means such as production of H_2_O_2_ (Calderón-Romero et al., 2018). In this sense, the crust of *Bacillus subtilis* is known to sustain heavy cross-link, most likely due to extensive disulfide bond formation between the cysteine-rich proteins encoded by the *cotVWXYZ* cluster composing this coat layer (Ursem et al., 2021; Zhang et al., 1993). In addition, CotY, is the protein of said layer that participates in the highest number of interactions with other coat proteins, even with itself alone (Krajčíková et al., 2017). Remarkably, it has been reported that heterologously produced CotY is capable of self-assembling into thermally stable structures in *E.coli* (and therefore, in absence of specialized sporulation machinery) that would only completely disaggregate in presence of strong reducing agents such as DTT at 99 °C for 20 minutes, likely due to the integrity conferred by such disulfide bonds (Jiang et al., 2015). Importantly, changes in the proteomic coat profile due to sporulation conditions presented several manners in which these cross-link reactions may be favoured in S_salt_ spores:

First, proteins involved in the enzymatic formation of all the three types of cross-link described in the *B. subtili*s spore coat, disulfide, dityrosine and ε-(γ-glutamyl) lysine bonds were more abundant in S_salt_ spores. For example, BdbD and SodA are either involved in or promote crosslinking reactions, such as disulfide bond formation between coat proteins (Henriques et al.,.1998; Krajčíková et al., 2017). Although the implication of the former protein is confirmed for adequate folding of secreted proteins containing cysteine, it was not needed for development of resistance in *Bacillus subtilis* spores to mild heat treatments of 80 °C for 15 minutes (Erlendsson et al., 2004). Whether it would provide protection against treatments of higher intensity is yet unknown. Conversely, deletion of SodA, is known to cause major coat structural alteration. In fact, it is known that the H_2_O_2_ produced by this superoxide dismutase can react with aminoacids involved in coat cross-linking such as cysteine and lysine (Finnegan et al., 2010; Ursem et al., 2021). And finally, it has been found to cross-link proteins such as CotG, although no significative effect on heat resistance of spores of strain AH1490 with no measurable superoxide dismutase activity was found (Henriques et al., 1998). In this context, it is important to note that the production of H_2_O_2_ by this enzyme family, and therefore, its ability to crosslink proteins, depends on the presence of superoxide radicals, which are more abundant under oxidative conditions, such as high salinity (Goosens et al., 2013; Hassan et al., 2020; Höper et al., 2006). In any case, the contribution of this proteins, especially the latter due to H_2_O_2_ production, by affecting cross-linking of proteins rich in these aminoacids, such as CotY is a tempting explanation that remains open for further investigation. In addition to this, the coat transglutaminase TgL known to favour cross-linking by ε-(γ-glutamyl) lysine bonds (Sanchez-Salas et al., 2011) was also present in higher abundance in S_salt_ spores, although the differences did not meet the cut-off criteria of FC > 2, potentially suggesting a more limited contribution to the observed phenotype.

Secondly, several proteins involved in various types of physical protein-protein interactions within the coat, including some that interact directly with CotY or self-interact, were also more abundant in S_salt_ (Table 2). For instance, increased presence of some of the proteins involved in polymerization control and scaffolding of the coat structure such as SpoVID and SafA may also favour a more robust coat structure (Costa et al., 2006; Ozin et al., 2000). In this sense, YabG cysteine protease may interact with both SpoVID and SafA with a similar mission to develop coat structure and resistance (Takamatsu et al., 2000). However, the essential role of CotY observed in the experiments supports the importance of proteins that either interact with it directly or are located in its immediate vicinity. Such could be the case for CotW, CotX, CotM and/or CgeA (Krajčíková et al., 2017; Liu et al., 2016) which showed an increase in their FC higher than 1.5 in S_salt_ (Table 2, underlined proteins). In this context, increased CotW may function as an anchor between the crust and the rest of the spore coat structure (Bartels et al., 2019). Similarly, the possibility that CotX contributes to the physical attachment of the crust to the rest of the coat was proposed by Shuster et al. (2019). Moreover, they also observed remarkable interactions among these proteins as location of CotY depended on CotW and codepended on CotX, while this latter was stabilized by CotW. Likewise, also demonstrated that CgeA localization dependend on both CotX and CotY. Importantly, CotX physical interaction with CotY was confirmed by single-molecule force spectroscopy (Liu et al., 2016). Although promising, to the best of our knowledge, the interactions of these proteins and the role of crust anchoring in spore heat resistance have yet to be examined.

Finally, there is evidence that proteins on the surface of spores may undergo spontaneous disulfide bond formation due to oxidative conditions (Richter et al., 2015). In fact, as previously mentioned, salinity can increase the oxidative conditions to which cells are exposed to (Hassan et al., 2020; Höper et al., 2006), which may help explain the vital role of a crust protein such as CotY in developing the increased resistance observed in S_salt_ spores. Furthermore, high oxidative conditions would explain the increased presence of redox homeostasis proteins in these spores as well, either as structural coat components or perhaps being “trapped” or adsorbed in the coat structure as is known to occur with heterologous proteins in *B. subtilis* (Huang et al., 2010; Isticato et al., 2013). In fact, other proteomic studies have also revealed an important presence of redox homeostasis proteins in the spore coat, such as Tpx or SodA located in the spore outer coat by Abhyankar et al. (2015). Its presence in the coat, supports a relevant role for regulation of the formation and/or function of this structure as suggested by other authors (Henriques et al., 2004). Increased abundance of redox proteins such as CotJC, Tpx, TrxA, AhpC, SodA, and YneT may protect against overoxidation of critical proteins or prevent damage in critical structures such as the IM. In fact, higher concentration of the putative catalase CotJC or the peroxidase Tpx in conjunction with TrxA could protect from exogenous H_2_O_2_ and, similarly, from that produced under increased oxidizing conditions, such as salinity, by other enzymes that also showed increased abundance in this population, like SodA. Regardless of their implication in this supposed regulatory system and how it is balanced, further studies will be needed to determine their exact contribution to resistance of free spores against these agents.

In conjunction, the higher oxidative environment provoked by salinity to which crust elements could be directly exposed to, the likely increased production of H_2_O_2_ by SodA under salinity conditions promoting ROS, and finally increased presence of elements able to form this kind of cross-link interactions directly with CotY in S_salt_ could promote structural resilience and changes in the properties of the coat. To date, however, no mechanism has been found about exactly how coat cross-linking influences wet heat resistance of bacterial spores. The coat itself presumably stabilizes the membrane during heat stress presumably by exerting pressure over the cortex and innermost layers (Kanaan et al., 2022; Setlow & Christie, 2023). In addition, the role of the crust in contributing to the tight packaging of the underlying layers is also supported by the observations made with TEM microscopy images of *cotXYZ* mutants (Freitas et al., 2020; Zhang et al., 1993). Expanding upon this, our results with fluorescence anisotropy of WT and Δ*cotY* spores indicated a substantial increase in the lipid mobility of Δ*cotY* spores. As a consequence, CotY or CotY related cross-linking, whether by one or multiple of the mechanisms described above, may help stabilize their membranes during heat lethal treatment by delaying damage accumulation to the IM and its constituents, believed to be the ultimate responsible target for spore inactivation by this technology (Coleman et al., 2007; Coleman & Setlow, 2009). This would be in accordance with our findings that S_salt_ spores would take damage more slowly as revealed by plating in selective media with NaCl (Freire et al., 2023) and also slower DPA release as well, as presented in this work. Further studies are needed to elucidate how exactly redox homeostasis, coat, and especially, crust modifications involving CotY increase the resistance of bacterial spores depending on the environmental conditions present during their formation.

## Conclusions

Spores formed at high salinity may increase their presence in the food chain due to the current tendency of salinization of important sporulation niches such as soil. Importantly, previous characterization found that spores formed under these conditions in the case of *B. subtilis* displayed increased wet heat resistance. In this work, we also observed increased resistance to hydrogen peroxide (H₂O₂), a disinfectant extensively employed in food processing environments, including dairy production facilities. Furthermore, the results presented in this manuscript showed that the increase in resistance to both wet heat and hydrogen peroxide was dependant on the crust morphogenetic protein CotY. Fluorescence anisotropy data with mutants lacking this protein showed its involvement in the stabilization of the IM during physiological conditions. And finally, was suggested to do so as well during lethal heat stress by monitoring DPA release at 95 °C. Remarkably, these spores formed at increased salinity, displayed increased abundance of coat redox homeostasis proteins such as Tpx and SodA, and many other coat components involved to promote, or establish cross-link reactions, including those with CotY. Therefore, this study remarks the relevance of the coat as a source of variability in spore composition and resistance due to increased salinity sporulation conditions and offers knowledge that will enhance our capability to understand resistance mechanism of bacterial spores, allowing the design of novel control strategies. For example, studying treatments aimed at weakening the coat integrity or disrupt disulfide bonds may offer a decrease in the intensity of heat treatments required to inactivate spores or improve biocide effectivity.

## Acknowledgements

This study was supported by a grant [PID2019-104712RA-I00] from MCIN/AEI/10.13039/501100011033 (to Elisa Gayán), and a P.h.D. scholarship from the Government of Aragón (to V.F.). Authors acknowledge labheads for providing access to institutional resources and funding that supported part of the research underlying this manuscript. The authors would also like to acknowledge the “Servicio General de Apoyo a la InvestigacionSAI” (University of Zaragoza) in developing the automatic colony counting system and Dr. C. Gross and the National BIO-Resource Project (NBRP) for providing us with the BKE genome-scale deletion library. Proteomic analyses were performed in the Proteomics Core Research Facility of Servicios Científico Técnicos del CIBA (IACS-Universidad de Zaragoza). We gratefully acknowledge the contributions of Santiago Condón † to this work. His insight and collaboration were invaluable.

## Contributions of authors

Conceptualization: VF, SC. Methodology: VF, IO, SC. Formal analysis: VF, SC. Investigation: VF, IO. Writing - original draft: VF. Writing - review & editing: VF, SC, IO.

## Notes

### Competing Interest Statement

The authors have declared no competing interest.

### Summary of Updates

Some mistakes were made in the Figures names in the text and also to improve following by the reader the figures were included in the main text

## References

Abdelshafy, A. M., Neetoo, H., & Al-Asmari, F. (2024). Antimicrobial Activity of Hydrogen Peroxide for Application in Food Safety and COVID-19 Mitigation: An Updated Review. Journal of Food Protection, 87(7), 100306. 10.1016/j.jfp.2024.100306

Abhyankar, W., Pandey, R., Ter Beek, A., Brul, S., de Koning, L. J., & de Koster, C. G. (2015). Reinforcement of *Bacillus subtilis* spores by cross-linking of outer coat proteins during maturation. Food Microbiology, 45, 54–62. 10.1016/j.fm.2014.03.007

Abhyankar, W. R., Kamphorst, K., Swarge, B. N., van Veen, H., van der Wel, N. N., Brul, S., de Koster, C. G., & de Koning, L. J. (2016). The Influence of Sporulation Conditions on the Spore Coat Protein Composition of Bacillus subtilis Spores. Frontiers in Microbiology, 7. https://www.frontiersin.org/articles/10.3389/fmicb.2016.01636

André, S., Vallaeys, T., & Planchon, S. (2017). Spore-forming bacteria responsible for food spoilage. Research in Microbiology, 168(4), 379–387. 10.1016/j.resmic.2016.10.003

Apetroaie-Constantin, C., Mikkola, R., Andersson, M. a., Teplova, V., Suominen, I., Johansson, T., & Salkinoja-Salonen, M. (2009). *Bacillus subtilis* and *B. mojavensis* strains connected to food poisoning produce the heat stable toxin amylosin. Journal of Applied Microbiology, 106(6), 1976–1985. 10.1111/j.1365-2672.2009.04167.x

Arnao, M. B., Acosta, M., del Río, J. A., & García-Cánovas, F. (1990). Inactivation of peroxidase by hydrogen peroxide and its protection by a reductant agent. Biochimica Et Biophysica Acta, 1038(1), 85–89. 10.1016/0167-4838(90)90014-7

Bartels, J., Blüher, A., López Castellanos, S., Richter, M., Günther, M., & Mascher, T. (2019). The *Bacillus subtilis* endospore crust: Protein interaction network, architecture and glycosylation state of a potential glycoprotein layer. Molecular Microbiology, 112(5), 1576– 1592. 10.1111/mmi.14381

Beaman, T. C., & Gerhardt, P. (1986). Heat resistance of bacterial spores correlated with protoplast dehydration, mineralization, and thermal adaptation. Applied and Environmental Microbiology, 52(6), 1242–1246. 10.1128/AEM.52.6.1242-1246.1986

Beaman, T. C., Greenamyre, J. T., Corner, T. R., Pankratz, H. S., & Gerhardt, P. (1982). Bacterial spore heat resistance correlated with water content, wet density, and protoplast/sporoplast volume ratio. Journal of Bacteriology, 150(2), 870–877.

Berendsen, E. M., Boekhorst, J., Kuipers, O. P., & Wells-Bennik, M. H. J. (2016). A mobile genetic element profoundly increases heat resistance of bacterial spores. The ISME Journal, 10(11), 2633–2642. 10.1038/ismej.2016.59

Berendsen, E. M., Koning, R. A., Boekhorst, J., de Jong, A., Kuipers, O. P., & Wells-Bennik, M. H. J. (2016). High-Level Heat Resistance of Spores of Bacillus amyloliquefaciens and Bacillus licheniformis Results from the Presence of a spoVA Operon in a Tn1546 Transposon.

*Frontiers in* *Microbiology*, 7. https://www.frontiersin.org/articles/10.3389/fmicb.2016.01912

Beskrovnaya, P., Sexton, D. L., Golmohammadzadeh, M., Hashimi, A., & Tocheva, E. I. (2021). Structural, Metabolic and Evolutionary Comparison of Bacterial Endospore and Exospore Formation. Frontiers in Microbiology, 12. 10.3389/fmicb.2021.630573

Bressuire-Isoard, C., Bornard, I., Henriques, A. O., Carlin, F., & Broussolle, V. (2016). Sporulation Temperature Reveals a Requirement for CotE in the Assembly of both the Coat and Exosporium Layers of *Bacillus cereus* Spores. Applied and Environmental Microbiology, 82(1), 232–243. 10.1128/AEM.02626-15

Bressuire-Isoard, C., Broussolle, V., & Carlin, F. (2018). Sporulation environment influences spore properties in *Bacillus*: Evidence and insights on underlying molecular and physiological mechanisms. FEMS Microbiology Reviews, 42(5), 614–626. 10.1093/femsre/fuy021

Burton, A. T., DeLoughery, A., Li, G.-W., & Kearns, D. B. (2019). Transcriptional Regulation and Mechanism of SigN (ZpdN), a pBS32-Encoded Sigma Factor in *Bacillus subtilis*. mBio, 10(5). 10.1128/mBio.01899-19

Calderón-Romero, P., Castro-Córdova, P., Reyes-Ramírez, R., Milano-Céspedes, M., Guerrero-Araya, E., Pizarro-Guajardo, M., Olguín-Araneda, V., Gil, F., & Paredes-Sabja, D. (2018). *Clostridium difficile* exosporium cysteine-rich proteins are essential for the morphogenesis of the exosporium layer, spore resistance, and affect *C. difficile* pathogenesis. PLoS Pathogens, 14(8), e1007199. 10.1371/journal.ppat.1007199

Carlin, F., Brillard, J., Broussolle, V., Clavel, T., Duport, C., Jobin, M., Guinebretière, M.-H., Auger, S., Sorokine, A., & Nguyen-Thé, C. (2010). Adaptation of *Bacillus cereus*, an ubiquitous worldwide-distributed foodborne pathogen, to a changing environment. Food Research International, 43(7), 1885–1894. 10.1016/j.foodres.2009.10.024

Chada, V. G. R., Sanstad, E. A., Wang, R., & Driks, A. (2003). Morphogenesis of *Bacillus* Spore Surfaces. Journal of Bacteriology, 185(21), 6255–6261. 10.1128/JB.185.21.6255-6261.2003

Chen, G., Driks, A., Tawfiq, K., Mallozzi, M., & Patil, S. (2010). *Bacillus anthracis* and *Bacillus subtilis* spore surface properties and transport. Colloids and Surfaces. B, Biointerfaces, 76(2), 512–518. 10.1016/j.colsurfb.2009.12.012

Clair, G., Esbelin, J., Malléa, S., Bornard, I., & Carlin, F. (2020). The spore coat is essential for *Bacillus subtilis* spore resistance to pulsed light, and pulsed light treatment eliminates some spore coat proteins. International Journal of Food Microbiology, 323(C), 108592. 10.1016/j.ijfoodmicro.2020.108592

Coleman, W. H., Chen, D., Li, Y., Cowan, A. E., & Setlow, P. (2007). How Moist Heat Kills Spores of *Bacillus subtilis*. Journal of Bacteriology, 189(23), 8458–8466. 10.1128/JB.01242-07

Coleman, W. h., & Setlow, P. (2009). Analysis of damage due to moist heat treatment of spores of *Bacillus subtilis*. Journal of Applied Microbiology, 106(5), 1600–1607. 10.1111/j.1365-2672.2008.04127.x

Condón, S., Arrizubieta, M. J., & Sala, F. J. (1993). Microbial heat resistance determinations by the multipoint system with the thermoresistometer TR-SC Improvement of this methodology. Journal of Microbiological Methods, 18(4), 357–366. 10.1016/0167-7012(93)90017-C

Cortezzo, D. e., & Setlow, P. (2005). Analysis of factors that influence the sensitivity of spores of *Bacillus subtilis* to DNA damaging chemicals. Journal of Applied Microbiology, 98(3), 606– 617. 10.1111/j.1365-2672.2004.02495.x

Costa, T., Isidro, A. L., Moran, C. P., & Henriques, A. O. (2006). Interaction between Coat Morphogenetic Proteins SafA and SpoVID. Journal of Bacteriology, 188(22), 7731–7741. 10.1128/jb.00761-06

Crow, A., Lewin, A., Hecht, O., Carlsson Möller, M., Moore, G. R., Hederstedt, L., & Le Brun, N. E. (2009). Crystal Structure and Biophysical Properties of *Bacillus subtilis* BdbD. The Journal of Biological Chemistry, 284(35), 23719–23733. 10.1074/jbc.M109.005785

Driks, A. (1999). *Bacillus subtilis* Spore Coat. Microbiology and Molecular Biology Reviews, 63(1), 1– 20.

Enger, K. S., Mitchell, J., Murali, B., Birdsell, D. N., Keim, P., Gurian, P. L., & Wagner, D. M. (2018). Evaluating the long-term persistence of *Bacillus* spores on common surfaces. Microbial Biotechnology, 11(6), 1048–1059. 10.1111/1751-7915.13267

Erlendsson, L. S., Möller, M., & Hederstedt, L. (2004). *Bacillus subtilis* StoA Is a Thiol-Disulfide Oxidoreductase Important for Spore Cortex Synthesis. Journal of Bacteriology, 186(18), 6230–6238. 10.1128/JB.186.18.6230-6238.2004

European Environment Agency (EEA). (2019). Soil moisture deficit. Retrieved from https://www.eea.europa.eu. (n.d.).

Fernandes, C. G., Martins, D., Hernandez, G., Sousa, A. L., Freitas, C., Tranfield, E. M., Cordeiro, T. N., Serrano, M., Moran, Charles. P., & Henriques, A. O. (2019). Temporal and spatial regulation of protein cross-linking by the pre-assembled substrates of a *Bacillus subtilis* spore coat transglutaminase. PLoS Genetics, 15(4), e1007912. 10.1371/journal.pgen.1007912

Fernandes, C. G., Moran, C. P., & Henriques, A. O. (2018). Autoregulation of SafA Assembly through Recruitment of a Protein Cross-Linking Enzyme. Journal of Bacteriology, 200(14), e00066–18. 10.1128/JB.00066-18

Finnegan, M., Linley, E., Denyer, S. P., McDonnell, G., Simons, C., & Maillard, J.-Y. (2010). Mode of action of hydrogen peroxide and other oxidizing agents: Differences between liquid and gas forms. Journal of Antimicrobial Chemotherapy, 65(10), 2108–2115. 10.1093/jac/dkq308

Flores, M. J., Duricy, K., Choudhary, S., Laue, M., & Popham, D. L. (2023). A Family of Spore Lipoproteins Stabilizes the Germination Apparatus by Altering Inner Spore Membrane Fluidity in *Bacillus subtilis* Spores. *Journal of Bacteriology*, e0014223. 10.1128/jb.00142-23

Freire, V., Condón, S., & Gayán, E. (2024). Impact of sporulation temperature on germination of *Bacillus subtilis* spores under optimal and adverse environmental conditions. Food Research International, 182, 114064. 10.1016/j.foodres.2024.114064

Freire, V., Condón, S., & Gayán, E. (2025). Sporulation at Reduced Water Activity Impairs Germination Kinetics of *Bacillus subtilis* Spores. Appl Environ Microbiol.

Freire, V., del Río, J., Gómara, P., Salvador, M., Condón, S., & Gayán, E. (2023). Comparative study on the impact of equally stressful environmental sporulation conditions on thermal inactivation kinetics of *B. subtilis* spores. International Journal of Food Microbiology, 405, 110349. 10.1016/j.ijfoodmicro.2023.110349

Freitas, C., Plannic, J., Isticato, R., Pelosi, A., Zilhão, R., Serrano, M., Baccigalupi, L., Ricca, E., Elsholz, A. K. W., Losick, R., & O. Henriques, A. (2020). A protein phosphorylation module patterns the *Bacillus subtilis* spore outer coat. Molecular Microbiology, 114(6), 934–951. 10.1111/mmi.14562

Geeson, N.A., Brandt, C.J., Thornes, J.B.,. (2003). Mediterranean desertification: A mosaic of processes and responses. (John Wiley&Sons).

Ghosh, A., Manton, J. D., Mustafa, A. R., Gupta, M., Ayuso-Garcia, A., Rees, E. J., & Christie, G. (2018). Proteins Encoded by the gerP Operon Are Localized to the Inner Coat in *Bacillus cereus* Spores and Are Dependent on GerPA and SafA for Assembly. Applied and Environmental Microbiology, 84(14), e00760–18. 10.1128/AEM.00760-18

Ghosh, S., Setlow, B., Wahome, P. G., Cowan, A. E., Plomp, M., Malkin, A. J., & Setlow, P. (2008). Characterization of Spores of *Bacillus subtilis* That Lack Most Coat Layers. Journal of Bacteriology, 190(20), 6741–6748. 10.1128/JB.00896-08

Goosens, V. J., Mars, R. A. T., Akeroyd, M., Vente, A., Dreisbach, A., Denham, E. L., Kouwen, T. R. H. M., van Rij, T., Olsthoorn, M., & van Dijl, J. M. (2013). Is proteomics a reliable tool to probe the oxidative folding of bacterial membrane proteins? Antioxidants & Redox Signaling, 18(10), 1159–1164. 10.1089/ars.2012.4664

Gould, G. W., & Hitchins, A. D. (1963). Sensitization of bacterial spores to lysozyme and to hydrogen peroxide with agents which rupture disulphide bonds. Journal of General Microbiology, 33, 413–423. 10.1099/00221287-33-3-413

Hassan, A. H. A., Alkhalifah, D. H. M., Al Yousef, S. A., Beemster, G. T. S., Mousa, A. S. M., Hozzein, W. N., & AbdElgawad, H. (2020). Salinity Stress Enhances the Antioxidant Capacity of *Bacillus* and *Planococcus* Species Isolated From Saline Lake Environment. Frontiers in Microbiology,1. 10.3389/fmicb.2020.561816

Henriques, A., Costa, T., Martins, L., & Zilhão, R. (2004). Functional architecture and assembly of the spore coat.

Henriques, A. O., Melsen, L. R., & Moran, C. P. (1998). Involvement of Superoxide Dismutase in Spore Coat Assembly in Bacillus subtilis. Journal of Bacteriology, 180(9), 2285–2291.

Höper, D., Bernhardt, J., & Hecker, M. (2006). Salt stress adaptation of *Bacillus subtilis*: A physiological proteomics approach. PROTEOMICS, 6(5), 1550–1562. 10.1002/pmic.200500197

Huang, J.-M., Hong, H. A., Van Tong, H., Hoang, T. H., Brisson, A., & Cutting, S. M. (2010). Mucosal delivery of antigens using adsorption to bacterial spores. Vaccine, 28(4), 1021–1030. 10.1016/j.vaccine.2009.10.127

Huang, Roseboom, W., Brul, S., & Kramer, G. (2023). Multi-omics analysis of Bacillus. Subtilis spores formed at different environmental temperatures reveal differences at the morphological and molecular level (p. 2023.06.22.546136). bioRxiv. 10.1101/2023.06.22.546136

Hullo, M. F., Moszer, I., Danchin, A., & Martin-Verstraete, I. (2001). CotA of *Bacillus subtilis* is a copper-dependent laccase. Journal of Bacteriology, 183(18), 5426–5430. 10.1128/JB.183.18.5426-5430.2001

Intergovernmental Panel on Climate Change (IPCC). (2022). Climate Change: Impacts, Adaptation and Vulnerability - Cross-Chapter Paper 3: Deserts, Semiarid Areas and Desertification. Retrieved from https://www.ipcc.ch/report/ar6/wg2/downloads/report/IPCC_AR6_WGII_FinalDraft_FullReport.pdf. (n.d.).

Isticato, R., Lanzilli, M., Petrillo, C., Donadio, G., Baccigalupi, L., & Ricca, E. (2020). *Bacillus subtilis* builds structurally and functionally different spores in response to the temperature of growth. Environmental Microbiology, 22(1), 170–182. 10.1111/1462-2920.14835

Isticato, R., Pelosi, A., Zilhão, R., Baccigalupi, L., Henriques, A. O., De Felice, M., & Ricca, E. (2008). CotC-CotU Heterodimerization during Assembly of the *Bacillus subtilis* Spore Coat. Journal of Bacteriology, 190(4), 1267–1275. 10.1128/JB.01425-07

Isticato, R., Sirec, T., Treppiccione, L., Maurano, F., De Felice, M., Rossi, M., & Ricca, E. (2013). Nonrecombinant display of the B subunit of the heat labile toxin of Escherichia coli on wild type and mutant spores of *Bacillus subtilis*. Microbial Cell Factories, 12, 98. 10.1186/1475-2859-12-98

Jamroskovic, J., Chromikova, Z., List, C., Bartova, B., Barak, I., & Bernier-Latmani, R. (2016). Variability in DPA and Calcium Content in the Spores of Clostridium Species. Frontiers in Microbiology, 7, 1791. 10.3389/fmicb.2016.01791

Jiang, S., Wan, Q., Krajcikova, D., Tang, J., Tzokov, S. B., Barak, I., & Bullough, P. A. (2015). Diverse supramolecular structures formed by self-assembling proteins of the *Bacillus subtilis* spore coat. Molecular Microbiology, 97(2), 347–359. 10.1111/mmi.13030

Kanaan, J., Murray, J., Higgins, R., Nana, M., DeMarco, A. M., Korza, G., & Setlow, P. (2022). Resistance properties and the role of the inner membrane and coat of *Bacillus subtilis* spores with extreme wet heat resistance. Journal of Applied Microbiology, 132(3), 2157– 2166. 10.1111/jam.15345

Khamidov, M., Ishchanov, J., Hamidov, A., Donmez, C., & Djumaboev, K. (2022). Assessment of Soil Salinity Changes under the Climate Change in the Khorezm Region, Uzbekistan. International Journal of Environmental Research and Public Health, 19(14), 8794. 10.3390/ijerph19148794

King, W. L., & Gould, G. W. (1969). Lysis of bacterial spores with hydrogen peroxide. The Journal of Applied Bacteriology, 32(4), 481–490. 10.1111/j.1365-2672.1969.tb01002.x

Klobutcher, L. A., Ragkousi, K., & Setlow, P. (2006). The *Bacillus subtilis* spore coat provides ‘eat resistance’ during phagocytic predation by the protozoan Tetrahymena thermophila. Proceedings of the National Academy of Sciences of the United States of America, 103(1), 165–170. 10.1073/pnas.0507121102

Knudsen, S. M., Cermak, N., Feijó Delgado, F., Setlow, B., Setlow, P., & Manalis, S. R. (2015). Water and Small-Molecule Permeation of Dormant *Bacillus subtilis* Spores. Journal of Bacteriology, 198(1), 168–177. 10.1128/JB.00435-15

Koo, B.-M., Kritikos, G., Farelli, J. D., Todor, H., Tong, K., Kimsey, H., Wapinski, I., Galardini, M., Cabal, A., Peters, J. M., Hachmann, A.-B., Rudner, D. Z., Allen, K. N., Typas, A., & Gross, C. A. (2017). Construction and Analysis of Two Genome-scale Deletion Libraries for *Bacillus subtilis*. Cell Systems, 4(3), 291–305.e7. 10.1016/j.cels.2016.12.013

Korza, G., DePratti, S., Fairchild, D., Wicander, J., Kanaan, J., Shames, H., Nichols, F. C., Cowan, A., Brul, S., & Setlow, P. (2023). Expression of the 2Duf protein in wild-type *Bacillus subtilis* spores stabilizes inner membrane proteins and increases spore resistance to wet heat and hydrogen peroxide. Journal of Applied Microbiology, 134(3), lxad040. 10.1093/jambio/lxad040

Krajčíková, D., Forgáč, V., Szabo, A., & Barák, I. (2017). Exploring the interaction network of the *Bacillus subtilis* outer coat and crust proteins. Microbiological Research, 204, 72–80. 10.1016/j.micres.2017.08.004

Leggett, M. J., Schwarz, J. S., Burke, P. A., McDonnell, G., Denyer, S. P., & Maillard, J.-Y. (2016). Mechanism of Sporicidal Activity for the Synergistic Combination of Peracetic Acid and Hydrogen Peroxide. Applied and Environmental Microbiology, 82(4), 1035–1039. 10.1128/AEM.03010-15

Liu, H., Krajcikova, D., Wang, N., Zhang, Z., Wang, H., Barak, I., & Tang, J. (2016). Forces and Kinetics of the *Bacillus subtilis* Spore Coat Proteins CotY and CotX Binding to CotE Inspected by Single Molecule Force Spectroscopy. The Journal of Physical Chemistry. B, 120(6), 1041–1047. 10.1021/acs.jpcb.5b11344

Liu, X. (2014). Membrane rigidity in vegetative cells and spores of Bacillus species [Rutgers University - Graduate School - New Brunswick]. 10.7282/T32Z13TR

Malyshev, D., Dahlberg, T., Wiklund, K., Andersson, P. O., Henriksson, S., & Andersson, M. (2021). Mode of Action of Disinfection Chemicals on the Bacterial Spore Structure and Their Raman Spectra. Analytical Chemistry, 93(6), 3146–3153. 10.1021/acs.analchem.0c04519

Marquis, R. E., & Bender, G. R. (1985). Mineralization and heat resistance of bacterial spores. Journal of Bacteriology, 161(2), 789–791. 10.1128/jb.161.2.789-791.1985

Mateo-Sagasta, & Burke. (2011). Agriculture and water quality interactions: A global overview. SOLAW Background Thematic Report—TR08.

McKenney, P. T., Driks, A., & Eichenberger, P. (2013). The *Bacillus subtilis* endospore: Assembly and functions of the multilayered coat. Nature Reviews Microbiology, 11(1), Article 1. 10.1038/nrmicro2921

Melly, E., Genest, P. c., Gilmore, M. e., Little, S., Popham, D. l., Driks, A., & Setlow, P. (2002). Analysis of the properties of spores of *Bacillus subtilis* prepared at different temperatures. Journal of Applied Microbiology, 92(6), 1105–1115. 10.1046/j.1365-2672.2002.01644.x

Monroe, A., & Setlow, P. (2006). Localization of the Transglutaminase Cross-Linking Sites in the *Bacillus subtilis* Spore Coat Protein GerQ. Journal of Bacteriology, 188(21), 7609–7616. 10.1128/JB.01116-06

Nguyen Thi Minh, H., Durand, A., Loison, P., Perrier-Cornet, J.-M., & Gervais, P. (2011). Effect of sporulation conditions on the resistance of *Bacillus subtilis* spores to heat and high pressure. Applied Microbiology and Biotechnology, 90(4), 1409–1417. 10.1007/s00253-011-3183-9

Nguyen Thi Minh, H., Guyot, S., Perrier-Cornet, J.-M., & Gervais, P. (2008). Effect of the osmotic conditions during sporulation on the subsequent resistance of bacterial spores. Applied Microbiology and Biotechnology, 80(1), 107–114. 10.1007/s00253-008-1519-x

Ozin, A. J., Henriques, A. O., Yi, H., & Moran, C. P. (2000). Morphogenetic Proteins SpoVID and SafA Form a Complex during Assembly of the *Bacillus subtilis* Spore Coat. Journal of Bacteriology, 182(7), 1828–1833.

Palop, A., Sala, F. J., & Condón, S. (1999). Heat Resistance of Native and Demineralized Spores of *Bacillus subtilis* Sporulated at Different Temperatures. Applied and Environmental Microbiology, 65(3), 1316–1319.

Ragkousi, K., & Setlow, P. (2004). Transglutaminase-Mediated Cross-Linking of GerQ in the Coats of *Bacillus subtilis* Spores. Journal of Bacteriology, 186(17), 5567–5575. 10.1128/JB.186.17.5567-5575.2004

Richter, A., Kim, W., Kim, J.-H., & Schumann, W. (2015). Disulfide Bonds of Proteins Displayed on Spores of *Bacillus subtilis* Can Occur Spontaneously. Current Microbiology, 71(1), 156–161. 10.1007/s00284-015-0839-1

Rose, R., Setlow, B., Monroe, A., Mallozzi, M., Driks, A., & Setlow, P. (2007). Comparison of the properties of *Bacillus subtilis* spores made in liquid or on agar plates. Journal of Applied Microbiology, 103(3), 691–699. 10.1111/j.1365-2672.2007.03297.x

Saggese, A., Di Gregorio Barletta, G., Vittoria, M., Donadio, G., Isticato, R., Baccigalupi, L., & Ricca, E. (2022). CotG Mediates Spore Surface Permeability in *Bacillus subtilis*. mBio, 13(6), e02760–22. 10.1128/mbio.02760-22

Sanchez-Salas, J.-L., Setlow, B., Zhang, P., Li, Y., & Setlow, P. (2011). Maturation of Released Spores Is Necessary for Acquisition of Full Spore Heat Resistance during *Bacillus subtilis* Sporulation. Applied and Environmental Microbiology, 77(19), 6746–6754. 10.1128/AEM.05031-11

Setlow, B., Atluri, S., Kitchel, R., Koziol-Dube, K., & Setlow, P. (2006). Role of Dipicolinic Acid in Resistance and Stability of Spores of *Bacillus subtilis* with or without DNA-Protective α/β- Type Small Acid-Soluble Proteins. Journal of Bacteriology, 188(11), 3740–3747. 10.1128/JB.00212-06

Setlow, P. (2014). Spore Resistance Properties. Microbiology Spectrum, 2(5). 10.1128/microbiolspec.TBS-0003-2012

Setlow, P., & Christie, G. (2023). New Thoughts on an Old Topic: Secrets of Bacterial Spore Resistance Slowly Being Revealed. Microbiology and Molecular Biology Reviews, 87(2), e00080–22. 10.1128/mmbr.00080-22

Setlow, P., & Johnson, E. A. (2019). Spores and Their Significance. In Food Microbiology (pp. 23–63). John Wiley & Sons, Ltd. 10.1128/9781555819972.ch2

Sevenich, R., & Mathys, A. (2018). Continuous Versus Discontinuous Ultra-High-Pressure Systems for Food Sterilization with Focus on Ultra-High-Pressure Homogenization and High-Pressure Thermal Sterilization: A Review. Comprehensive Reviews in Food Science and Food Safety, 17(3), 646–662. 10.1111/1541-4337.12348

Shin, S.-Y., Calvisi, E. G., Beaman, T. C., Pankratz, H. S., Gerhardt, P., & Marquis, R. E. (1994). Microscopic and Thermal Characterization of Hydrogen Peroxide Killing and Lysis of Spores and Protection by Transition Metal Ions, Chelators, and Antioxidants. Applied and Environmental Microbiology, 60(9), 3192–3197.

*Soil threats in Europe: Status, methods, drivers and effects on ecosystem services—ESDAC - European Commission*. (n.d.). Retrieved 27 May 2025, from https://esdac.jrc.ec.europa.eu/content/soil-threats-europe-status-methods-drivers-andeffects-ecosystem-services

Sunde, E. P., Setlow, P., Hederstedt, L., & Halle, B. (2009). The physical state of water in bacterial spores. Proceedings of the National Academy of Sciences of the United States of America, 106(46), 19334–19339. 10.1073/pnas.0908712106

Takamatsu, H., Imamura, A., Kodama, T., Asai, K., Ogasawara, N., & Watabe, K. (2000). The yabG gene of *Bacillus subtilis* encodes a sporulation specific protease which is involved in the processing of several spore coat proteins. FEMS Microbiology Letters, 192(1), 33–38. 10.1111/j.1574-6968.2000.tb09355.x

Tu, Z., Setlow, P., Brul, S., & Kramer, G. (2021). Molecular Physiological Characterization of a High Heat Resistant Spore Forming *Bacillus subtilis* Food Isolate. Microorganisms, 9(3), 667. 10.3390/microorganisms9030667

Tyanova, S., Temu, T., Sinitcyn, P., Carlson, A., Hein, M. Y., Geiger, T., Mann, M., & Cox, J. (2016). The Perseus computational platform for comprehensive analysis of (prote)omics data. Nature Methods, 13(9), 731–740. 10.1038/nmeth.3901

Ursem, R., Swarge, B., Abhyankar, W. R., Buncherd, H., de Koning, L. J., Setlow, P., Brul, S., & Kramer, G. (2021b). Identification of Native Cross-Links in *Bacillus subtilis* Spore Coat Proteins. Journal of Proteome Research, 20(3), 1809–1816. 10.1021/acs.jproteome.1c00025

Voss, D., & Montville, T. J. (2014). 1,6-Diphenyl-1,3,5-hexatrine as a reporter of inner spore membrane fluidity in Bacillus subtilis and Alicyclobacillus acidoterrestris. Journal of Microbiological Methods, 96, 101–103. 10.1016/j.mimet.2013.11.009

Wijman, J. G. E., de Leeuw, P. P. L. A., Moezelaar, R., Zwietering, M. H., & Abee, T. (2007). Air-liquid interface biofilms of *Bacillus cereus*: Formation, sporulation, and dispersion. Applied and Environmental Microbiology, 73(5), 1481–1488. 10.1128/AEM.01781-06

Zhang, J., Fitz-James, P. C., & Aronson, A. I. (1993). Cloning and characterization of a cluster of genes encoding polypeptides present in the insoluble fraction of the spore coat of *Bacillus subtilis*. Journal of Bacteriology, 175(12), 3757–3766.

Zhyvoloup, A., Yu, B. Y. K., Baković, J., Davis-Lunn, M., Tossounian, M.-A., Thomas, N., Tsuchiya, Y., Peak-Chew, S. Y., Wigneshweraraj, S., Filonenko, V., Skehel, M., Setlow, P., & Gout, I. (2020). Analysis of disulphide bond linkage between CoA and protein cysteine thiols during sporulation and in spores of *Bacillus* species. FEMS Microbiology Letters, 367(23), fnaa174. 10.1093/femsle/fnaa174

