## Supplemental figures 1 and 2 for "High-salinity sporulation in *Bacillus subtilis* results in coat dependant enhanced resistance to both wet heat and hydrogen peroxide"

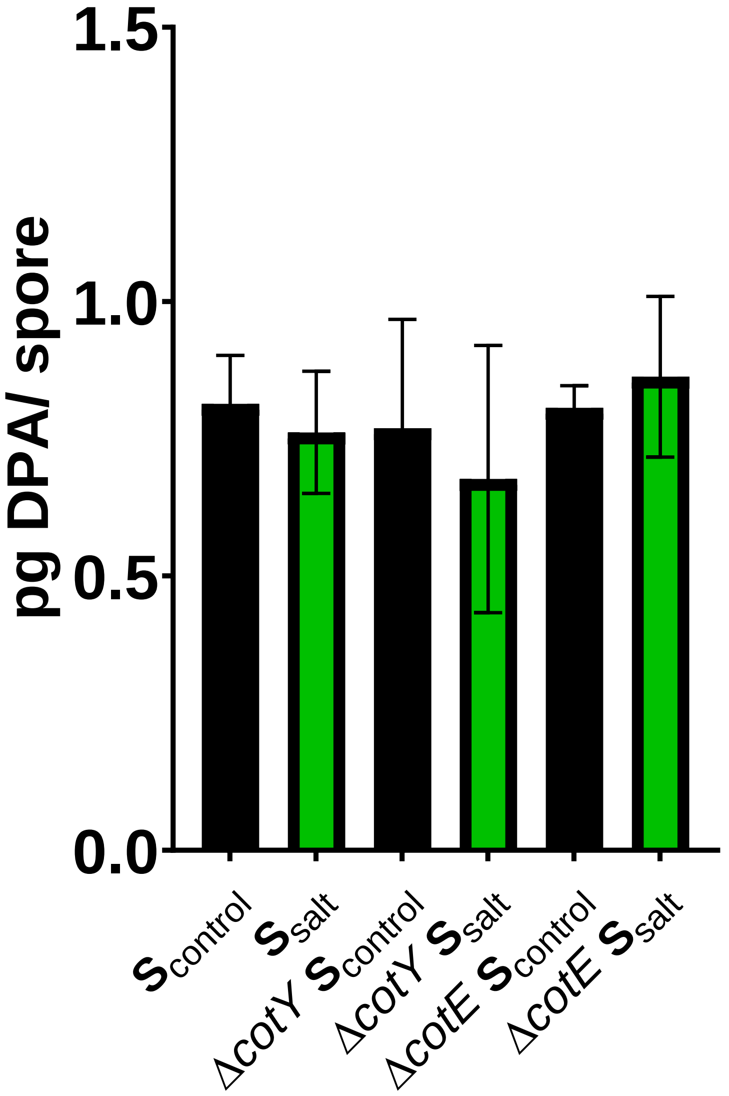


**Figure S1:** DPA content of WT and morphogenetic coat deficient strains of *B. subtilis* 168 sporulated under both optimal (black) and under high salinity conditions (green).

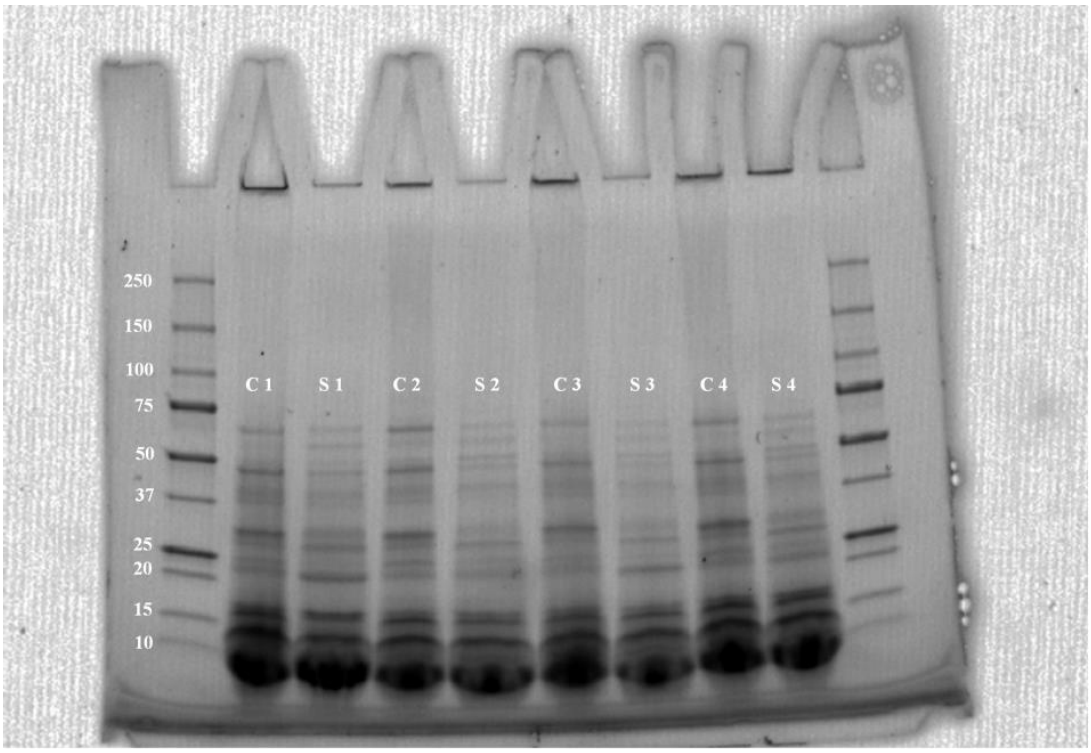


**Figure S2.** SDS-PAGE 2D electrophoresis of coat protein extracts. Lanes 1 and 10: molecular-weight markers (kDa). Lanes 2, 4, 6, and 8: $S_{control}$ samples (C). Lanes 3, 5, 7, and 9: $S_{salt}$ samples (S).
